# Gain-of-function CRISPR screens identify tumor-promoting genes conferring melanoma cell plasticity and therapy-resistance

**DOI:** 10.1101/2020.07.08.193102

**Authors:** Arthur Gautron, Laura Bachelot, Anaïs M. Quéméner, Sébastien Corre, Marc Aubry, Florian Rambow, Anaïs Paris, Nina Tardif, Héloïse M. Leclair, Cédric Coulouarn, Jean-Christophe Marine, Marie-Dominique Galibert, David Gilot

**Author notes:** These authors contributed equally to this work (Anaïs M. Quéméner, Sébastien Corre, Marc Aubry). Correspondence or, Corresponding Author: David Gilot, 2 avenue Pr Léon Bernard, 35043 Rennes, FRANCE., or Marie-Dominique Galibert, 2 avenue Pr Léon Bernard, 35043 Rennes, FRANCE.

## Abstract

Most genetic alterations that drive melanoma development and resistance to targeted therapy have been uncovered. In contrast, and despite their increasingly recognized contribution, little is known about the non-genetic mechanisms that drive these processes. Here, we performed *in vivo* gain-of-function CRISPR screens and identified *SMAD3*, *BIRC3* and *SLC9A5* as key actors of BRAFi-resistance and these genes promote the tumor growth capability of persister cells. We show that their expression levels increase during acquisition of BRAFi-resistance, and remain high in persister cells and during relapse. The upregulation of the SMAD3 transcriptional activity (SMAD3-signature) promotes a mesenchymal-like phenotype and BRAFi-resistance by acting as an upstream transcriptional regulator of potent BRAFi-resistance genes such as EGFR and AXL. This SMAD3-signature predicts resistance to both current melanoma therapies in different cohorts. Critically, chemical inhibition of SMAD3 may constitute amenable target for melanoma since it efficiently abrogates persister cells survival. Interestingly, decrease of SMAD3 activity can also be reached by inhibiting the aryl hydrocarbon receptor (AhR), another druggable transcription factor governing SMAD3 expression level. Our work expands our understanding of the biology of persister cells and highlight novel drug vulnerabilities that can be exploited to develop long-lasting antimelanoma therapies.

## INTRODUCTION

Identifying molecular cancer drivers is critical for precision oncology. Last year, the cancer genome atlas (TCGA) identified 299 driver genes by focusing on point mutations and small indels across 33 cancer types ^1^. It represents the most comprehensive effort thus far to identify cancer driver mutations. Complementary studies are required to elucidate the role of copy-number variations, genomic fusions and methylation events in the 33 TCGA projects.

Moreover, there is increasing evidence that non-genetic reprogramming leading to cancer cell dedifferentiation, stemness, invasiveness also contribute to tumor growth and therapy-resistance ^2, 3^. Thus, deciphering the signaling pathways that drive such processes may also lead to innovative cancer therapies. Recent gene expression quantifications performed at single cell level by single cell RNA sequencing (scRNA-Seq) demonstrated that cancer cells operate a dedifferentiation process, for instance in glioblastoma and melanoma ^4–6^, promoting tumor growth, stemness and therapy-resistance. This ‘onco-dedifferentiation’ seems to be independent of *de novo* mutations and could offer new targets/strategies to cure cancer. However, these scRNA-Seq studies are mainly descriptive; the tumor growth capability of each gene/RNA is not yet investigated at the genome-scale. Such functional analyses are nowadays feasible using clustered regularly interspaced short palindromic repeats (CRISPR)-Cas9 screens ^7^. The majority of the CRISPR-Cas9 screens is based on the invalidation of coding genes but modulation of gene expression is reachable with the CRISPR-Cas9 synergistic activation mediator (SAM) approach ^8^. It corresponds to an engineered protein complex for the transcriptional activation of endogenous genes. Importantly, SAM can further be combined with a human genome-wide library to activate all known coding isoforms from the RefSeq database (23,430 isoforms) for gain-of-function screening without *a priori*. To date, CRISPR screens are mainly performed *in vitro* using cell lines or primary cultures ^9^. A pan-cancer CRISPR-Cas9 knock-out screen was performed *in vitro* (324 human cancer cell lines from 30 cancer types) to identify essential genes for cancer cell fitness (defined as genes required for cell growth or viability) and to prioritize candidates for cancer therapeutics ^10^. However, because the contribution of the tumor environment in tumor growth is increasingly recognized, it seems important to perform such screens in the relevant patho-physiological context and, for instance, take advantage of animal models.

We selected cutaneous melanoma as a paradigm since novel therapeutic strategies are critically needed ^3^. Targeted therapies such as BRAF inhibitors (BRAFi) initially showed great promise in patients with BRAF(V600)-mutated metastatic melanoma. Unfortunately, the vast majority of patients that initially respond to these drugs, almost inevitably develop resistance. Although combination therapies (BRAF and MEK inhibitors) enhance the response and delay relapse, the overall survival remains unsatisfactory highlighting the need of new therapeutic targets ^11^.

The mechanisms underlying resistance are numerous and probably not mutually exclusive ^12–14^. Resistance can be driven by a small pre-existing subpopulation, harboring specific genetic alterations that confer them with resistance to the inhibitors ^15^. Such alterations may also occur *de novo*, during treatment ^12^. In addition, there is increased evidence that non-genetic reprogramming may confer drug tolerant and/or resistant phenotypes to melanoma cells ^6, 16–20^. Earlier works demonstrated that phenotype switching from a proliferative to an invasive/mesenchymal-like state is also likely to contribute to therapy resistance ^21–25^. Paradoxically, MITF-induced differentiation into a slow cycling, pigment-producing state was also reported to confer tolerance to BRAFi ^25, 26^. It therefore seems that various drug tolerant subpopulations can emerge under therapeutic pressure and that these cells can provide a pool from which resistance develops. Targeting these populations of persister cells is therefore crucial to achieve effective personalized therapies ^27^.

Here, we performed unbiased screens to identify genes promoting tumor growth from persister cells and conferring resistance to BRAFi using CRISPR-Cas9 SAM methodology. We demonstrate that, in addition to promote melanoma development, *SMAD3*, *BIRC3* and *SLC9A5* also support relapse since they promote both BRAFi-resistance and tumor growth capability of persister cells. Their expression levels correlated with BRAFi-resistance and relapse. Consequently, their inhibition strongly reduced the number of persister cells. Moreover, we demonstrate that the transcription factor AhR governs *SMAD3* expression levels and in turn SMAD3 drives the expression of a set of genes associated with BRAFi-resistance and mesenchymal phenotype. These experiments identify integrated AhR-SMAD3 signaling as a key driver of melanoma growth and relapse, pointing to a new therapeutic vulnerability in melanoma.

## RESULTS

### Identification of Tumor-Promoting Genes by *in vivo* Gain-of-Function CRISPR Screen

Since the tumor environment influences, at least in part, the tumor growth capability of cancer cells, we performed an *in vivo* genome-wide CRISPR-Cas9 SAM screen to identify *in vivo* tumor-promoting genes, defined as genes whose expression support tumor growth (in contrast to driver genes bearing a driver mutation such as BRAF (V600E)). To select the most appropriate cellular model, we classified melanoma biopsies from The Cancer Genome Atlas (TCGA) cohort (n = 458) in function of differentiation states according to the most recent melanoma profiling data (Fig. 1A) ^17^. As anticipated, the vast majority of these tumors, which are almost all drug naive, exhibited a differentiated profile (89%; melanocytic and transitory). We selected the 501Mel cell line since i) these cells display a melanocytic differentiation state as the majority of diagnosticated melanoma, ii) they harbor the BRAF(V600E) mutation as ∼50% of cutaneous melanoma, iii) they are highly sensitive to BRAFi with an IC_50_ value of 0.45 μM to vemurafenib [PLX4032] ^18, 28^ and importantly iv) they are unable to generate tumor in immunodeficient mice ^29^. This latter characteristic may allow to identify tumor-promoting genes.

**Figure 1.**
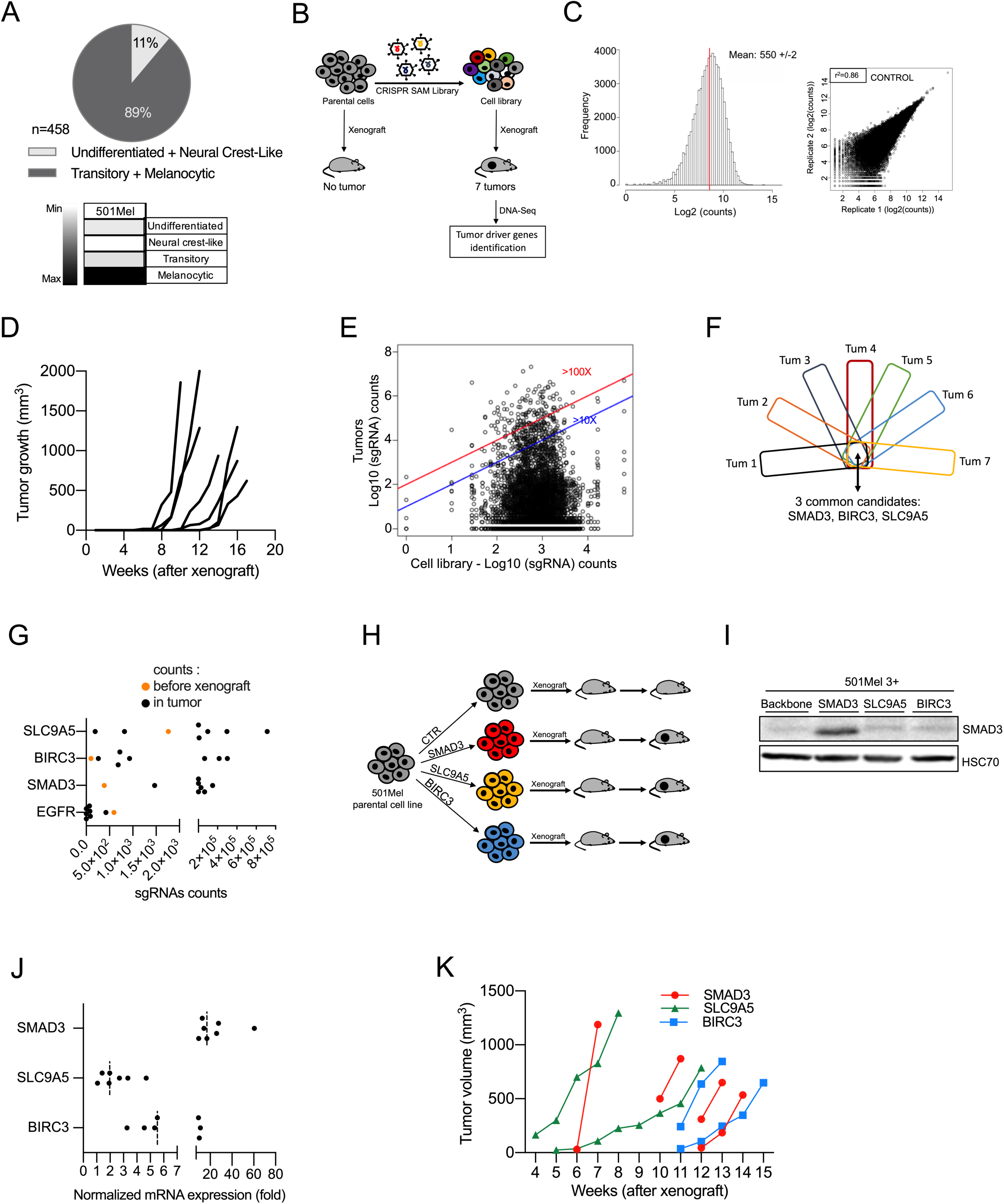
Identification of Tumor-Promoting Genes by *in vivo* Gain-of-Function CRISPR Screen. (A) Determination of differentiation states of skin cutaneous melanoma (SKCM) biopsies from the TCGA cohort (n=458) according to ^17^. The vast majority of these tumors exhibited a differentiated profile (89%; melanocytic and transitory states). The others display a dedifferentiated profile (11%; neural crest-like cells and undifferentiated states). Human melanoma 501Mel cell line is classified as differentiated melanoma cells, according to ^17^ and selected for the CRISPR screens. (B) Workflow depicting the *in vivo* CRISPR-SAM screen to identify tumor-promoting genes. Parental cells and cell library were xenografted on nude mice (3×10^6^cells/mouse, n=6 and n=10, respectively) and tumor growth was monitored during 5 months. Seven tumors were collected and analyzed by DNA-Seq to identify the sgRNAs. (C) left: sgRNAs distribution in the cell library. sgRNA mean reaches 550+/-2. Right: Inter-replicate correlation between the two biological replicates was determined using the Pearson correlation coefficient r. sgRNAs(log2(counts)). (D) Tumor growth curves for the 7 tumors arising from the CRISPR-SAM-engineered cells xenografted in nude mice as detailed in Fig. 1B. (No tumor for the parental cell line (501Mel cells)). (E) Distribution of sgRNAs in cell library and in the 7 tumors (log_10_(sgRNAs counts)). Blue and red lines indicated the enrichment ≥10 fold or ≥100 fold (tumors vs cell library (*in vitro*)). (F) From the seven tumors, the top hundred genes (enriched) have been selected and common genes are SMAD3, BIRC3 and SLC9A5. (G) sgRNA counts in tumors *versus* in cell library (respectively, black and orange points). Each black point corresponds to one tumor. (H) Workflow depicting the validation step : 501Mel cells overexpressing *SMAD3*, *BIRC3* or *SLC9A5* (obtained by CRISPR-SAM) were xenografted on nude mice and tumor volume was monitored using caliper. 3×10^6^ cells/mouse. n=7, 6 and 6 mice, respectively. (I) SMAD3 expression levels in 501Mel cells overexpressing the cofactors for CRISPR-SAM approach and the control sgRNA (501Mel 3+ backbone) or the SMAD3 sgRNA. SLC9A5 and BIRC3 sgRNAs are used as controls to show the specificity of the SMAD3 overexpression. HSC70 serves as loading control. (J) *SMAD3*, *BIRC3* and *SLC9A5* mRNA expression levels in melanoma cell lines described in H and I. n=7 independent biological experiments. (K) Tumor growth curves from 501Mel cells overexpressing *SMAD3*, *BIRC3* or *SLC9A5.* Western blot results are representative of at least two independent experiments. Source data are available in Table S10 and unprocessed original blots are shown in Fig. S6. See also Fig. S1.

To generate the CRISPR-SAM cell library, we modified the 501Mel cells, to express constitutively defective-Cas9 and the required cofactors for CRISPR-SAM technology ^8^. These engineered cells were infected with the single-guide RNA (sgRNA) lentivirus library that contained at least three different guides per coding gene ^8^ (Fig. 1B). The infection was performed at a multiplicity of infection (MOI) of 0.2 ensuring that only one guide is expressed per infected cell. Infected cells were positively selected using antibiotic selection during 7 days. By DNA-sequencing, we observed a normal distribution of the sgRNAs in two cell library replicates (Fig. 1C). Only 78 sgRNAs are not detected in our cell library, which validate our protocol and the cell library (>70,100 sgRNAs were detected) (Table S1). The quality of the two cell library replicates was evaluated by estimating the distribution of the guides (Fig. 1C, right panel). Thus, all the quality controls were proper to identify *in vivo* tumor-promoting genes.

The cell library (30×10^6^ cells) was fractionated and subcutaneously xenografted in 10 nude mice (3×10^6^ cells/mouse) and tumor growth was monitored using caliper over a 5 months period (Fig. 1D and table S2). As previously demonstrated ^29^, we confirmed that parental 501Mel are unable to form tumors in nude mice (n=6). In contrast, seven tumors were obtained from the CRISPR-engineered cells (Fig. 1D). The nature of the sgRNAs, their abundance and occurrence across these 7 tumors were determined by DNA-Seq (Fig. 1E, and Table S3).

By comparing the most represented genes (sgRNAs) in each tumor (Tum), we identified 3 common genes (Fig.1F). An enrichment of SMAD3, BIRC3 and SLC9A5 sgRNAs was found in tumors when compared to their starting abundance (cell library; orange points) (Fig.1G), in contrast to EGFR sgRNAs. Thirty-six other genes were recurrently retrieved in the tumors but not in all (Table S3). Interestingly, *YAP1*, which has already been identified as melanoma growth-promoting gene was also found (Table S3). This supports the robustness of the screen ^16, 22, 30^. The majority of the tumor-promoting genes (Table S4) identified here are not considered as genes required for cell growth or viability (except the essential genes *YAP1, SLC25A41* and *TGIF1*) ^10^, and are not frequently altered in melanoma (Fig S1A). These results suggest that a high expression level of these genes is sufficient to promote melanoma tumor growth. Moreover, the transforming growth factor (TGF)-β pathway seemed well-represented among the tumor-promoting genes (Table S4).

Next, we examined *in vivo* the ability of these tumor-promoting genes to promote tumor growth by xenografting cell populations overexpressing the sgRNAs individually. We focused on the top 3 tumor-promoting genes; *SMAD3*, *BIRC3* and *SCL9A5* (Fig. 1H-K). We generated three new CRISPR-engineered cell lines and we evaluated the overexpression levels of these genes by western-blot experiments (SMAD3) and RT-qPCR (SMAD3, BIRC3 & SLC9A5) (Fig. 1I & 1J). Finally, we confirmed that they independently foster tumor development (Fig. 1K). Altogether, our results demonstrated that *in vivo* CRISPR-SAM screen identifies new tumor-promoting genes, which may constitute amenable target for melanoma.

### Genome-wide CRISPR Activation Screen Identifies BRAFi-Resistance Genes in Cutaneous Melanoma

BRAFi provoke tumor shrinkage in the vast majority of patients with BRAF(V600)-mutated metastatic melanoma but resistance almost inevitably occurs ^3^. To examine the potential role of the above identified tumor-promoting genes (Fig. 1) in BRAFi-resistance and relapse, we performed an *in cellulo* screen using the same cell library in the presence of BRAFi (Fig. 2). Briefly, the CRISPR-SAM 501Mel cell library (40×10^6^ cells) was treated for 14 days with BRAFi (2 μM), using either the BRAFi used in clinical practice (vemurafenib), the next generation inhibitor that is still under investigation in clinical trials (PLX8394), or the solvent (dimethyl sulfoxide (DMSO)) as control. This procedure allows for the enrichment of sgRNAs (genes) conferring resistance. The nature of the sgRNA present in the resistant population and their abundance was determined by DNA-Seq (Fig. 2B). The best hit was the Epidermal growth factor receptor gene (*EGFR*), a well-known BRAFi-resistance gene ^31, 32^. By examining the enrichment of sgRNAs targeting *EGFR* promoter (Fig. 2B), we decided to retain genes with at least two sgRNAs among the enriched sgRNAs present in BRAFi-exposed cells (with a false discovery rate, FDR <0.05) (Table S6) since sometimes one of the three sgRNAs designed per gene is not detected or not enriched as observed for *EGFR* (Fig. 2C). A recent publication confirmed that sgRNAs are not all functional in CRISPRa libraries and it could be interesting to increase the number of sgRNAs per target and to cover more TSS per gene ^33^.

**Figure 2.**
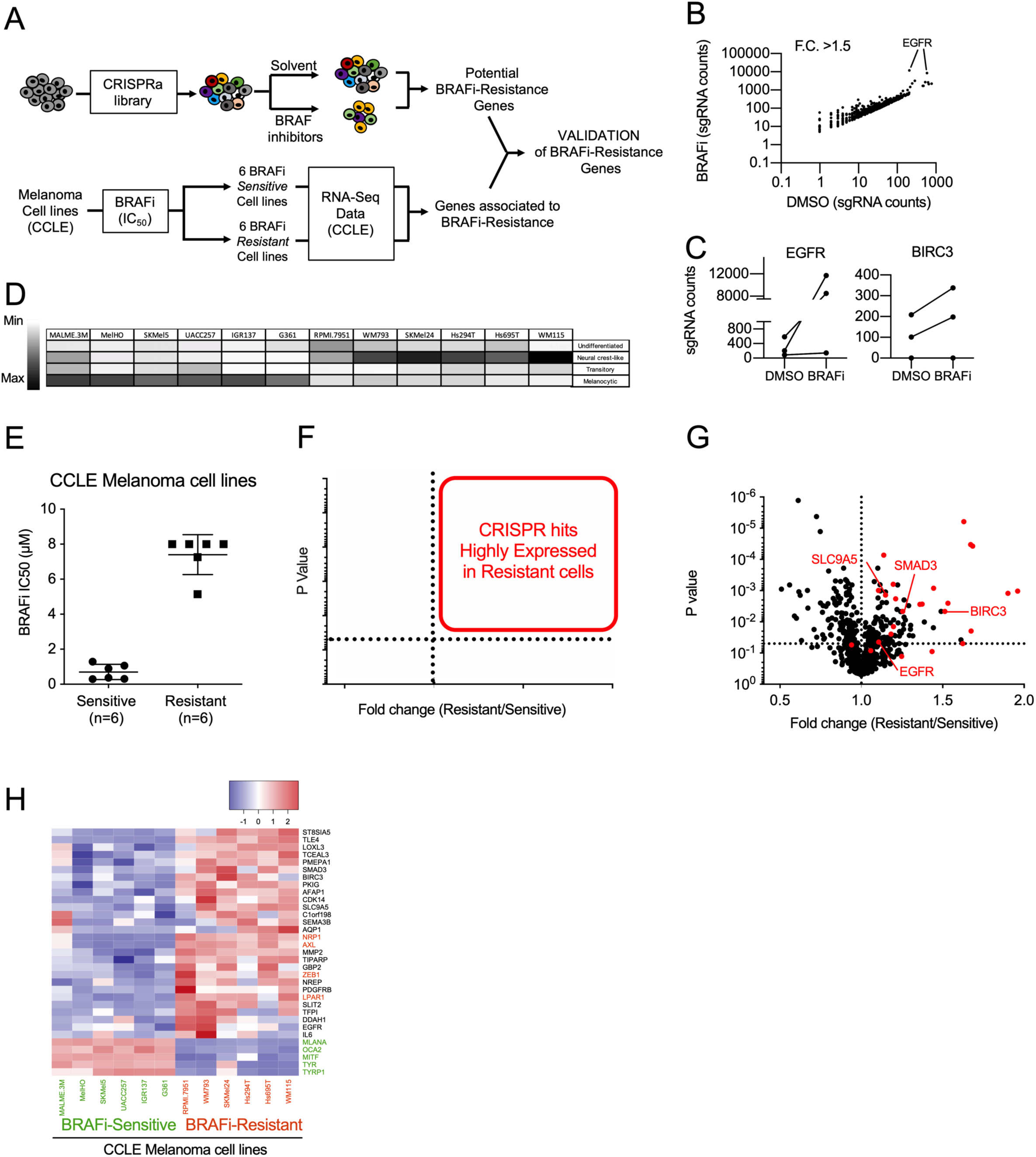
Genome-wide CRISPR Activation Screen Identifies BRAFi-Resistance Genes in Cutaneous Melanoma. (A) CRISPR-SAM workflow. Cell library was exposed to DMSO (solvent of BRAFi) or BRAFi during 14 days. 40×10^6^ cells per arm. Experiments were done in duplicate. (B) Plot showing sgRNAs detected in BRAFi-resistant cells *versus* in control cells (cell library exposed to DMSO) with a fold change >1.5. Raw data are available in Table S5. (C) sgRNAs counts in BRAFi-resistant cells *versus* in DMSO-exposed cells for EGFR and BIRC3. (D) Determination of differentiation states of 12 human BRAF(V600) melanoma cell lines from CCLE according to ^17^. (E) BRAF(V600) melanoma cell lines from CCLE were distributed in two groups (Sensitive and Resistant) according to their BRAFi IC_50_ (μM, half maximal inhibitory concentrations) ^34^. (F) Volcano plot explaining the identification of BRAFi-resistant genes found both in the screen and enriched in BRAFi-resistant cell lines. (G) Volcano plot showing the expression levels of BRAFi-resistant genes (identified in 501Mel by CRISPR-SAM screen) in 12 human melanoma cell lines. The fold change corresponds to the ratio of expression levels found in resistant and sensitive cell lines. In red: selected genes. *EGFR* is considered as positive control and *SMAD3*, *BIRC3* and *SLC9A5*; selected as favorite genes for the next steps. *SMAD3*, *BIRC3*, *SLC9A5* and *AFAP1* are BRAFi-resistance genes and potent tumor-promoting genes (Fig. 1). Raw data are available in Table S5. (H) Heatmap recapitulating the expression levels of the selected hits (red dots in Fig. 2G) in BRAFi-resistant and -sensitive cell lines. Markers of differentiation (*MITF*, *MLANA*, *OCA2*, *TYRP1*, *TYR*). In blue: resistance genes already published (*NRP1*, *AXL*, *ZEB1*, *LPAR1*).

Apart sgRNAs targeting the *EGFR* promoter, the sgRNAs enrichment in BRAFi-exposed cells were unexpectedly weak (Tables S5 and S6). Thus, to select the best BRAFi-resistance genes, we examined the gene expression levels of these potential BRAFi-resistance genes identified by CRSIPR screen in 12 melanoma cell lines (Fig. 2D and 2E). We postulated that BRAFi-resistance genes are highly expressed in BRAFi-resistant cells (n=6) as already demonstrated by other approaches for *NRP1*, *AXL*, *NGFR*. To this end, we confronted CRISPR-SAM candidates (identified in 501Mel cells) to gene expression data from six melanoma cell lines that were highly resistant to BRAFi according to the Cancer Cell Lines Encyclopedia (CCLE) *versus* six sensitive cell lines ^34^ (Fig. 2E). We next focused on candidate genes which were both enriched in CRISPR screen and highly expressed in the majority of BRAFi-resistance cell lines (Fig. 2F and 2G). To better evaluate the validity of our candidate genes identified using this workflow, we added well-established and validated BRAFi-resistant genes (*NRP1*, *AXL*, *ZEB1* and *LPAR1)* ^8, 25, 35^ and five genes associated with melanoma cell differentiation (*MITF*, *OCA2*, *MLANA*, *TYR*, and *TYRP1*) ^36^. All BRAFi-resistant cell lines presented a dedifferentiated profile as anticipated. Importantly, our candidate genes including *SMAD3, SLC9A5* and *BIRC3* (Fig. 1H, black color) displayed a similar expression profile than observed for the well-established and validated BRAFi-resistant genes (Fig. 2H, red color), strongly suggesting that these genes may also confer BRAFi-resistance. *EGFR* and platelet-derived growth factor receptor (*PDGFR)-β* showed high expression only in a few BRAFi-resistant cell lines, as previously observed in patients ^31^.

Together, our results confirmed the robustness of the functional *in cellulo* CRISPR-SAM screen.

### Validation of BRAFi SAM-selected Resistance Genes

To evaluate the contribution of the CRISPR-SAM-selected genes in BRAFi resistance, we examined the transcriptome of the differentiated cell line M229 (melanocytic) at different stages during acquisition of resistance (Fig. 3A) ^13^. As described, cell lines were exposed to chronic exposure to BRAFi and analysed at different days of treatment (P: parental cells, 2D: two days of treatment, DTP: drug tolerant persister cells, DTPP: drug tolerant proliferating persister cells, SDR: single-drug resistant cell). We observed a sequential upregulation in the expression of BRAFi SAM-selected genes: a group of genes (*MMP2*, *SEMA3B*, *BIRC3*, *TIPARP*, etc) being expressed earlier than a 2^nd^ group (*IL-6*, *EGFR*, *AFAP1*, etc). The majority of the candidates were overexpressed while the cells exhibited resistance to a single BRAFi agent (single drug resistance, SDR). Comparable results were obtained with the M238 melanoma cell line (Fig. 3B). Importantly, we found that combining the BRAFi with MEKi (DDR) led to comparable up-regulation of the BRAFi SAM-selected genes than observed in cells exposed to BRAFi alone (SDR) (Fig. 3C). These results indicate that a common gene expression program can confer resistance to inhibitors of MAPK pathway as previously reported ^18, 37^.

**Figure 3.**
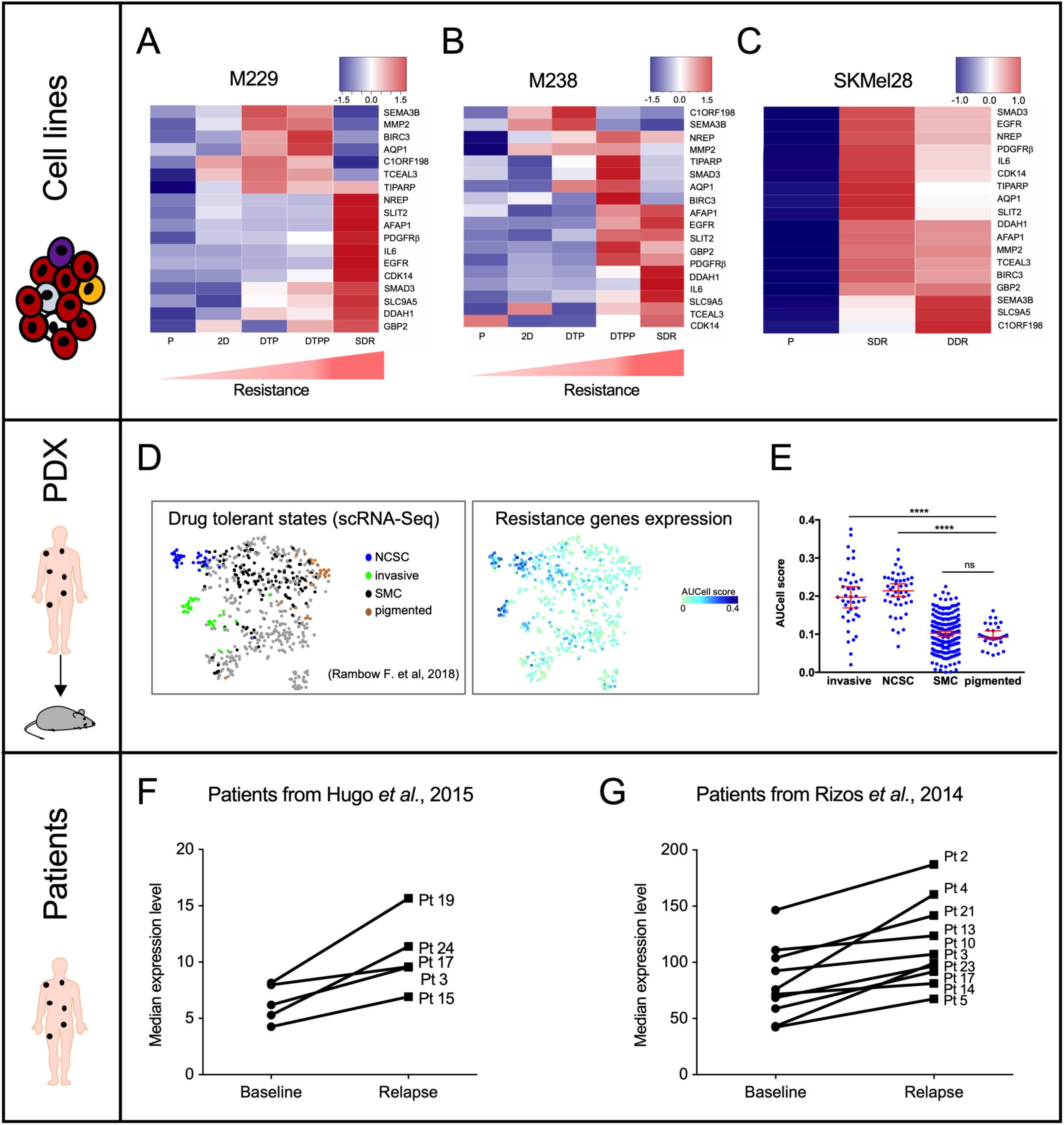
Validation of BRAFi SAM-selected Resistance Genes. (A) Expression levels of our hits in M229 melanoma cells during the BRAFi-resistance acquisition. P: parental cells, 2D: two days of treatment, DTP: drug tolerant persister cells, DTPP: drug tolerant proliferating persister cells, SDR: single-drug resistant cells (BRAFi). (B) Expression levels of our hits in M238 melanoma cells during the BRAFi-resistance acquisition. (C) Expression levels of our hits in parental, single drug resistant (SDR, BRAFi) or dual drug resistant (DDR, BRAFi+MEKi) SKMel28 melanoma cell lines. (D) T-distributed Stochastic Neighbor Embedding (t-SNE) plot showing the 4 drug tolerant states (NCSC (neural crest stem cells), invasive, SMC (starved-like melanoma cells) and pigmented cells) according to single cell RNA-seq (scRNA-Seq) performed in PDX model exposed to BRAFi+MEKi ^6^. Our BRAFi-resistance genes are mainly expressed in NCSC and invasive cells. (E) AUCell score for each drug tolerant states. ****p<0.0001, Mann-Whitney test. (F) Expression level of our genes (median) in cutaneous melanoma biopsies before the BRAFi-treatment (baseline) and during the relapse. Cohort from ^16^. (G) Expression level of our genes (median) in cutaneous melanoma biopsies before the BRAFi-treatment and during the relapse. Cohort from ^66^. See also Fig. S2.

Having shown that combination of BRAFi and MEKi promotes sequential up-regulation of BRAFi SAM-selected genes, we monitored their expression levels in an *in vivo* preclinical patient-derived xenograft (PDX) model ^6^. Using scRNA-Seq, our collaborators reported the presence of dedifferentiated drug tolerant cells exhibiting a neural crest stem cell (NCSC) and invasive profiles at minimal residual disease isolated from the MEL006 PDX model ^6^. Here, we showed that the BRAFi SAM-selected genes were highly and selectively expressed in both of these cell populations (Fig. 3D and 3E). Moreover, *in silico* analyses of EGFR-expressing cells sorted from melanoma tumors displayed comparatively high expression levels of the BRAFi SAM-selected genes (Fig. S2A & S2B). Importantly, these EGFR-positive cells are able to proliferate in the presence of BRAFi and generate BRAFi-resistant colonies ^32^. In addition, high expression levels of the BRAFi SAM-selected genes have been found in invasive cells when compared to proliferative melanoma cell lines (Fig. S2C). Together, these data confirm the upregulated expression of BRAFi SAM-selected genes in cells shown to contribute to relapse, further supporting their involvement in establishing drug tolerant and/or resistant phenotypes *in vivo*.

To evaluate the clinical relevance of the SAM-selected BRAFi-resistance genes, we compared their expression levels (median) in two independent BRAFi drug naive/drug resistant patient cohorts (Fig. 3F and 3G and Fig. S2D and S2E). The expression levels of the selected resistant candidate genes increased during relapse in the majority of drug resistant patients. Notably, none of these genes have been implicated in recurrent gene-amplification events that are sometimes identified in drug naive lesions (cBioPortal, TCGA) ^38^ (Fig. S2F-S2H). These data indicate that the increase in expression of the SAM-selected genes in BRAFi-resistant cells is likely associated with a (non-genetic) dedifferentiation process of melanoma cells induced by the therapy. Together, these *in vitro* and *in vivo* gene expression analyses strongly support a BRAFi-resistance function for the SAM-selected genes.

### BRAFi-Resistance Genes Promote Tumor Growth

Long-term effect of BRAFi is reduced by the ability of persister cells to resist to BRAFi but also to promote the tumor growth (relapse). Thus, we investigated the capability of the BRAFi-persister cells (obtained from the *in vitro* CRISPR-screen, Fig. 2) to promote tumor growth into immune-deficient mice. The subset of BRAFi-resistant/persister cells (Fig. 4A) were engrafted (36×10^6^ cells, 3×10^6^/mouse) and tumor growth was monitored (Fig. 4B).

**Figure 4.**
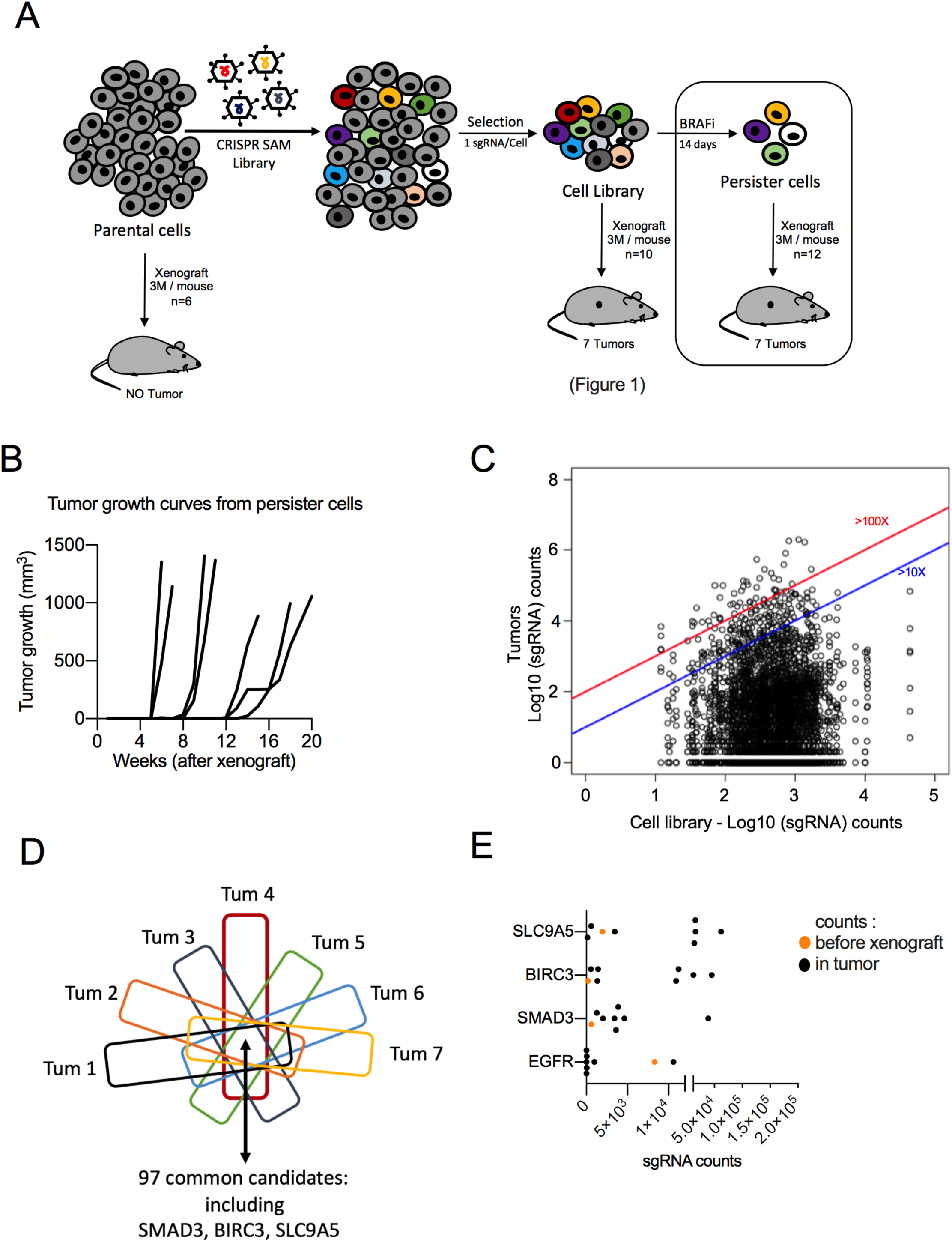
BRAFi-Resistance Genes Promote Tumor Growth. (A) Workflow to identify genes involved in tumor growth from BRAFi-persister cells (36×10^6^ cells xenografted in 12 mice: 3×10^6^ persister cells/mouse). (B) Tumor growth curves from BRAFi-persister cells (monitored during 5 months). (C) Distribution of sgRNAs in BRAFi-resistant cells (before xenograft) and in the 7 tumors emerging from the resistant/persister cells (log_10_(sgRNAs counts)). Blue and red lines indicated the enrichment ≥10 fold or ≥100 fold (tumors *versus* BRAFi-resistant cells (*in vitro*)). Raw data are available in Table S7. (D) From the seven tumors arising from persister cells, the common genes (enriched) have been extracted. Ninety-seven genes including SMAD3, BIRC3 and SLC9A5 are detected in all these 7 tumors. Raw data are available in Table S7. (E) sgRNAs counts in tumors *versus* sgRNA detected in BRAFi-resistant cell library (*in vitro*) (respectively, black and orange points) for selected candidates. Each black point corresponds to one tumor. *EGFR* was the most potent BRAFi-resistant gene (Fig.2). *SMAD3*, *BIRC3* and *SLC9A5* were identified as hits in the three screens (Fig.1, 2 and 4).

These BRAFi-persister cells formed tumors, in contrast to parental 501Mel cells ^29^. We determined the nature and abundance of sgRNAs present in each emerging tumor (Fig. 4C and Table S7) and we identified 97 genes (sgRNAs) detected in all tumors, including *SMAD3*, *BIRC3* and *SLC9A5* (Fig. 4D). We looked for the enrichment of these sgRNAs in each tumor developed from persister cells (Fig. 4E). *EGFR* sgRNA was not frequently enriched in tumors (as previously described for human melanoma tumors ^31, 32, 39^) in contrast to *SMAD3*, *BIRC3*, and *SLC9A5* (Fig. 4E). It is important to note that we already identified these 3 genes as tumor-promoting genes and BRAFi-resistant genes in previous figures. Thus, these 3 genes are potential interesting targets for antimelanoma therapy.

### Functional Validation of Genes Involved in BRAFi-Resistance and Relapse

We focused on the transcription factor SMAD3 as a model gene for monitoring BRAFi-resistance and relapse due to its critical function downstream of the TGFβ pathway. Although this pathway is known to promote melanoma phenotype switching/dedifferentiation ^31^, to support melanoma growth ^40^ and metastasis, little is known about the role of SMAD3 in melanoma biology and as a modulator of resistance to targeted therapy.

We confirmed that *SMAD3* mRNA is highly expressed in dedifferentiated cells (Fig. 5A-5C) and in BRAFi-resistant cells (Fig. 2H).

**Figure 5.**
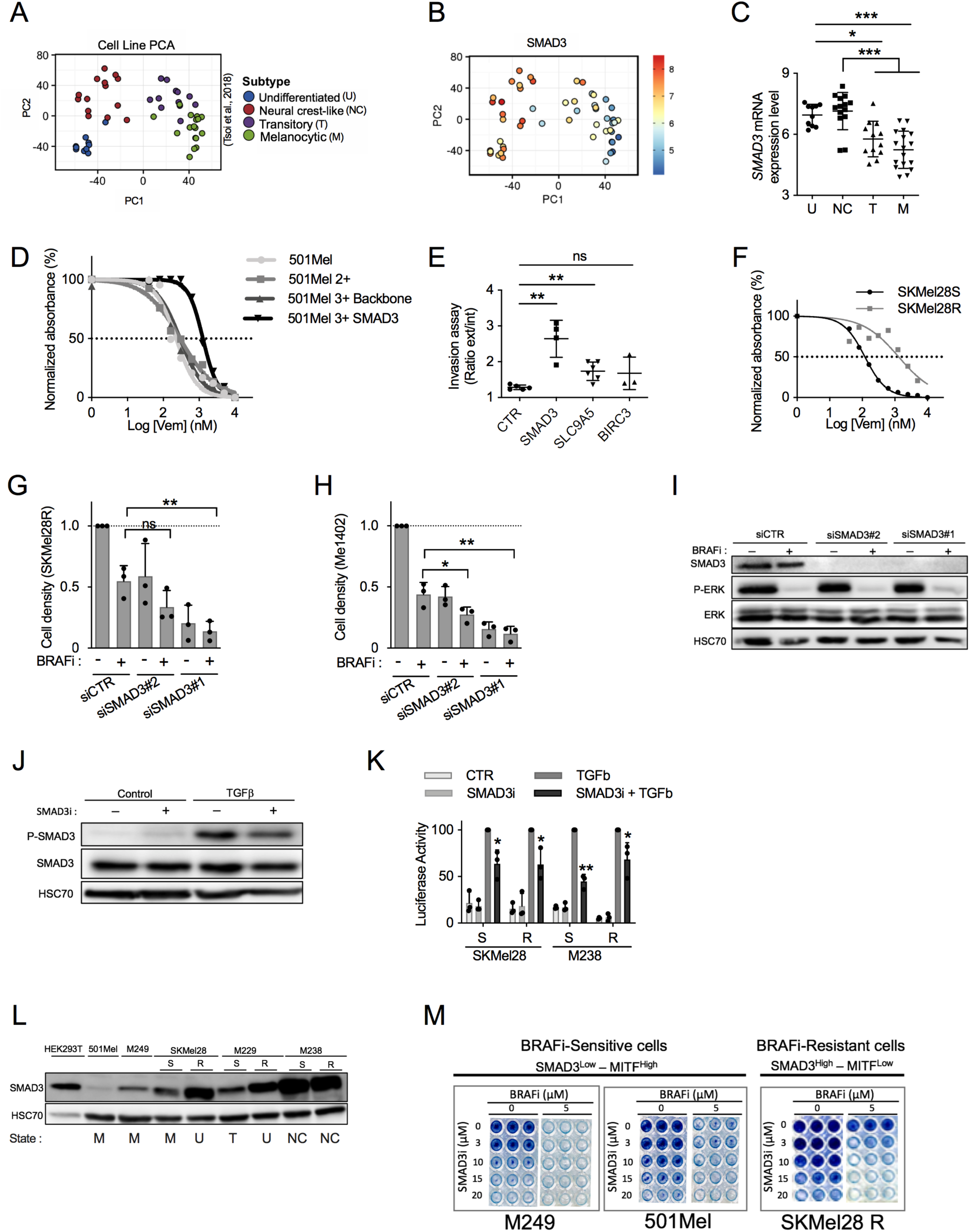
Functional Validation of Genes Involved in BRAFi-Resistance and Relapse. (A) PCA analysis of melanoma cell lines in function of their dedifferentiation states (generated by the webtool http://systems.crump.ucla.edu/dediff/index.php). (B) SMAD3 expression increases with melanoma cell dedifferentiation. PCA analysis of *SMAD3* expression in melanoma cell lines in function of their dedifferentiation states. Scale: red color corresponds to a high *SMAD3* expression level. (C) *SMAD3* expression in these four subtypes of melanoma cells (U, undifferentiated; N, neural crest-like; T, transitory; M, melanocytic). Number in each group: U = 10, N = 14, T = 12, M =17. Whiskers reflect mean of expression with range. Bilateral Student test (with non-equivalent variances) *: p<0.05, ***p: <0.001. (D) SMAD3 gain-of-function increases BRAFi-resistance. Determination of BRAFi half maximal inhibitory concentrations (IC_50_ values) for 501Mel cell lines (Log[Vem] (nM)). Parental 501Mel cells (n=5), 501Mel cells expressing dCas9 and HSF1-p65-MS2 (named here 501Mel 2+, n=5), the 501Mel 2+ cells expressing a control guide (backbone, n=5) and the 501Mel 2+cells overexpressing SMAD3 (n=3 biologically independent experiments). A representative experiment has been chosen. (E) SMAD3 and SLC9A5 gain-of-function increases invasion capability of melanoma cells. Invasion assays for engineered cells lines (<u>melanoma spheroids)</u>: CTR; 501Mel 2+, SMAD3; 501Mel 2+cells overexpressing SMAD3. Two other cell lines overexpressing SLC9A5 or BIRC3 have been tested. Explanation for ratio calculation is detailed in Fig. S3. Results obtained from two biologically independent experiments. (n=5, 4, 6 and 3 spheroids, respectively). Whiskers reflect mean of values with s.d.. Bilateral Student test (with non-equivalent variances) **: p<0.01. (F) Characterization of SKMel28 sensitive-and resistant-cell lines (SKMel28S and SKMel28R). Determination of BRAFi half maximal inhibitory concentrations (IC_50_ values) (Log[Vem] (nM)). A representative experiment has been chosen among two experiments. (G) *SMAD3* depletion (siRNA#1 & #2) decreased cell density and increased BRAFi effect (vemurafenib) on BRAFi-resistant cells (SKMel28R). CTR for non-targeting siRNA. DMSO for dimethylsulfoxide (solvent of vemurafenib; BRAFi). n**=**3 biologically independent experiments. Each histogram represents the mean ± s.d. ; Bilateral Student test (with non-equivalent variances) **: p<0.01. (H) *SMAD3* depletion (siRNA#1 & #2) decreased cell density and increased BRAFi effect (vemurafenib, BRAFi) on BRAFi-resistant cells (Me1402). CTR for non-targeting siRNA. DMSO for dimethylsulfoxide (solvent of vemurafenib). n**=**3 biologically independent experiments. Each histogram represents the mean ± s.d. ; Bilateral Student test (with non-equivalent variances) *: p<0.05, **: p<0.01. (I) Validation of SMAD3 knock-down by western blot experiments in Me1402 cells exposed or not to Vemurafenib (BRAFi 5μM, 2 days). Cells were exposed to BRAFi 24hr after siRNA transfection. Vemurafenib inhibitory effect on mutated BRAF was evaluated by analysing the phospho-ERK1/2 levels. ERK and HSC70 serves as loading control. (J) Validation of SMAD3 inhibitor (SIS3, SMAD3i). Effect of SMAD3i (SIS3, 10μM) on the level of phospho-SMAD3 Ser423/425 in response to TGFβ (2ng/mL, 1hr) (or solvent: HCl 4mM + Bovine serum albumin 1mg/mL) in 501Mel cells overexpressing SMAD3 using CRISPR-SAM. Serum starved cells (500,000 per well) were pretreated with SMAD3i 10μM (or control solvent) during 2hr before TGFβ addition. SMAD3 and HSC70 serve as loading control for western blot experiments. (K) Inhibitory effect of SMAD3i (SIS3) on the SMAD-responsive luciferase activity. Vector encodes the Firefly luciferase reporter gene under the control of a minimal (m)CMV promoter and tandem repeats of the SMAD Binding Element (SBE). Cells (10,000 per well) were pretreated with SAMD3i 10μM or control solvent during 1.5hr, and next cells were exposed to TGFβ 10ng/mL (or solvent: HCl 4mM + Bovine serum albumin 1mg/mL) for 6hr. n**=**3 biologically independent experiments. Each histogram represents the mean ± s.d. ; Bilateral Student test (with non-equivalent variances); *: p<0.05, **: p<0.01 (L) SMAD3 expression levels increase in BRAFi-resistant cell lines and in dedifferentiated cells. Four subtypes of melanoma cells (U, undifferentiated; N, neural crest-like; T, transitory; M, melanocytic) have been used. 501Mel and M249 cells are melanocytic cells in contrast to dedifferentiated BRAFi-resistant cells (R). Three couples of melanoma cell lines (Sensitive (S) and R) have been used to illustrate the SMAD3 increase in BRAFi-resistant cell lines. HEK293T cells are used as control (kidney). HSC70 serves as loading control for western blot experiments. (M) The combo (SMAD3i and BRAFi) eradicated the persister cells. Chemical inhibition of SMAD3 by SIS3 (SMAD3i) restored vemurafenib effect (5μM, 4 days) on BRAFi-resistant cells (SKMel28R). [SMAD3i]; 0, 3, 10, 15 or 20 μM during 4 days. BRAFi-sensitive cells (501Mel & M249) are SMAD3^low^ and MITF^high^ and are highly sensitive to single treatment (BRAFi (5μM)) in contrast to BRAFi-resistant cells. Representative pictures of n**=**2 biologically independent experiments. Western blot results are representative of at least two independent experiments. Source data are available in Table S10 and unprocessed original blots are shown in Fig. S6. See also Fig. S3.

To reinforce the role of SMAD3 in BRAFi-resistance, we showed that gain-of-function of SMAD3 significantly increases the BRAFi-resistance of melanoma cells when compared to different control cells (parental 501Mel cells and the CRISPR-engineered cells : 501Mel cells expressing dCas9 and HSF1-p65-MS2 (named here 501Mel 2+) and the 501Mel 2+cells expressing a control guide (named 3+ backbone) (Fig. 5D). In addition, we showed that gain-of-function of SMAD3 also promotes the three-dimensional (3D) tumor spheroid invasion capability of melanoma cells. Similar results were obtained for SLC9A5 (Fig. 5E & Fig. S3A). These results strongly suggest that a high expression level of *SMAD3* confers BRAFi-resistance and invasion capability in melanoma cells.

Thus, we investigated if SMAD3 impairment re-sensitizes cells to BRAF inhibitor, using small-interfering RNA (siRNA) (Fig. 5F-I). To select an accurate *in vitro* model, we examined the expression level of *SMAD3* using the gene expression webtool developed by Graeber’s team ^17^ (n=53 cell lines) (Fig. 5A). We found that *SMAD3* expression levels are higher in the dedifferentiated cells than in the differentiated ones (Fig. 5A and 5B). As the dedifferentiation status correlated with BRAFi resistance, we selected two BRAFi-resistant cell lines, SKMel28 BRAFi-resistant cells (SKMel28R, Fig. 5F, and 5G) ^16^ and Me1402 melanoma cells (Fig. 5H). In contrast to the SKMel28R, the resistance of which were created by chronic exposure to non-lethal doses of BRAFi, Me1402 cells are intrinsically resistant.

Surprisingly, the single SMAD3 depletion decreased the cell density in a similar magnitude than BRAFi treatment in these BRAFi-resistant cells (Fig. 5G and 5H). The *SMAD3* depletion did not modify the ERK pathway (Fig. 5I) in contrast to the BRAFi. The combo (*SMAD3* depletion and BRAFi (5μM)) efficiently reduced the number of resistant/persister cells in these two cell lines, suggesting that SMAD3 is an interesting target to limit resistance to BRAFi. Similar results were obtained by targeting *BIRC3*, *EGFR*, *IL6* or *AQP1* (Fig. S3B-S3G).

To transfer this strategy (SMAD3 inhibition + BRAFi) into clinic, we looked for an efficient inhibitor of SMAD3. We identified the chemical inhibitor SIS3 (SMAD3 inhibitor, SMAD3i) ^41^. Firstly, we validated the inhibitory efficiency of SMAD3i in melanoma cells since it strongly decreased the levels of phospho-SMAD3 Ser423/425 induced by TGFβ (Fig. 5J) in accordance with previous studies ^41, 42^. Moreover, we demonstrated that SMAD3i reduced the transcriptional activity of SMAD3 in response to TGFβ exposure in 4 melanoma cell lines (Fig 5K). To know if the combination (SMAD3i+BRAFi) could be broadly used to eradicate persister cells emerging in response to BRAFi treatment, we selected three melanoma cell lines in function of *SMAD3* expression levels (Fig. 5L). Differentiated cells (501Mel and M249; SMAD3^low^) are highly sensitive to BRAFi (decrease of cell density: >90% at 5μM BRAFi, 4 days) and the combo eradicated the rare persister cells with a low dose of SMAD3i. Dedifferentiated cells (SKMel28R, SMAD3^high^) are highly resistant to BRAFi (decrease of cell density: ∼50% at 5μM BRAFi, 4 days) and the combo eradicated the persister cells (Fig. 5M). Together, these results identified *SMAD3* as an amenable target to limit resistance to BRAFi and tumor growth.

### The Transcription Factor AhR Drives *SMAD3* Expression

Having shown that SMAD3 expression mediates BRAFi resistance and tumor growth, we explored the transcriptional program promoting its expression in BRAFi-resistant melanoma cells. We recently reported that the Aryl hydrocarbon Receptor (AhR), a ligand-dependent transcription factor is an upstream central node regulating the expression of BRAFi resistance genes and melanoma dedifferentiation ^18^.

We postulated that AhR may govern *SMAD3* expression in BRAFi-resistant cells. We identified three putative canonical binding sites for AhR (XRE for xenobiotic responsive element) in the proximal promoter of *SMAD3* (Fig. 6A) in accordance to chromatin immunoprecipitation coupled to massively parallel DNA sequencing data (AhR ChIP-Seq) showing AhR binding on SMAD3 promoter ^43, 44^. To demonstrate the *SMAD3* induction by AhR, 501Mel cells were exposed to the most potent and well-known AhR ligands (TCDD for 2,3,5,7-tetrachlorodibenzodioxine; ITE for 2-(1*H*-Indol-3-ylcarbonyl)-4-thiazolecarboxylic acid methyl ester). These AhR ligands increased *SMAD3* expression, in an AhR-dependent manner (Fig. 6B). Comparable results were obtained with a canonical target gene of AhR; the TCCD-induced poly(ADP ribose) polymerase gene (*TIPARP*), supporting the role of AhR in regulating *SMAD3* expression (Fig. 6C). The need of an activated-AhR promoting *SMAD3* expression was further confirmed by the use of an AhR-antagonist (CH-223191) in SKMel28 cells (Fig. 6D). Long-term chemical inhibition of AhR activity reduced *SMAD3* and *TIPARP* expression levels. In accordance with these results, *SMAD3* expression levels decreased in AhR KO SKMel28 cells (Fig. 6E).

**Figure 6.**
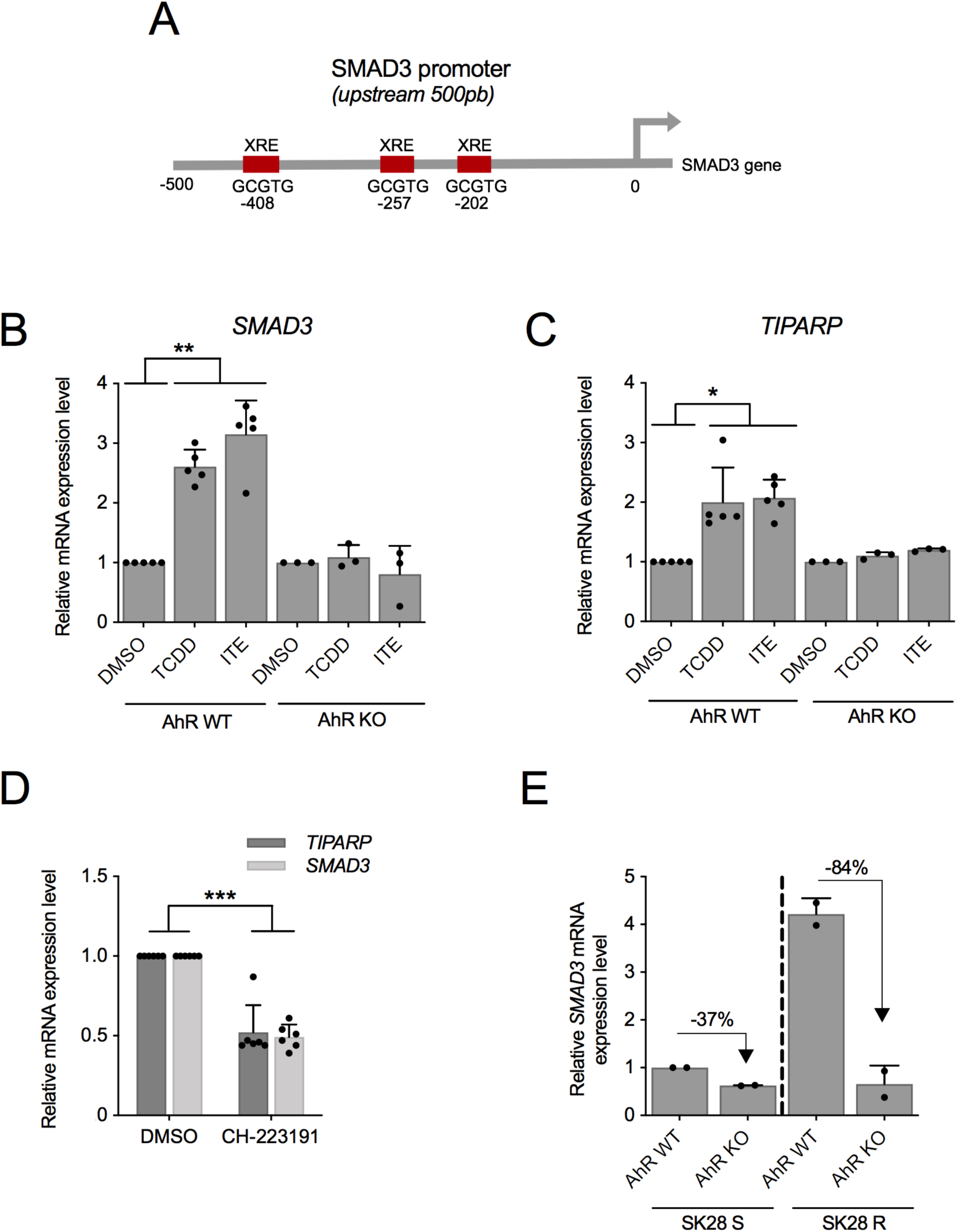
The Transcription Factor AhR Drives *SMAD3* Expression. (A) AhR binding sites (xenobiotic responsive element (XRE); GCGTG) in human *SMAD3* proximal promoter (−500_+1 bp). (B) AhR activation by exogenous and endogenous ligands promotes *SMAD3* induction. 501Mel cells AhR wild-type or knock-out have been exposed to exogenous and endogenous AhR ligands; TCDD (5nM) or ITE (10μM) or the solvent (DMSO) during 10 days. n**=**5 biologically independent experiments for AhR WT cells and n=3 for AhR KO cells. Each histogram represents the mean ± s.d.; Bilateral Student test (with non-equivalent variances): **\*\***, p**<**0.01. (C) AhR activation by exogenous and endogenous AhR ligands promotes *TIPARP* induction. 501Mel cells have been treated as described in B. n**=**5 biologically independent experiments for AhR WT cells and n=3 for AhR KO cells. Each histogram represents the mean ± s.d.; Bilateral Student test (with non-equivalent variances): **\***, p**<**0.05. (D) AhR antagonist (CH-223191) reduces *SMAD3* and *TIPARP* expression levels. SKMel28 cells (AhR wild-type) have been exposed to CH-223191 (5μM) or the solvent (DMSO) during 7 days. n**=**6 biologically independent experiments. Each histogram represents the mean ± s.d.; Bilateral Student test (with non-equivalent variances): **\*\*\*:** p**<**0.001. (E) Loss of AhR reduces *SMAD3* expression levels. SMAD3 expression has been investigated in SKMel28 cells AhR wild-type (WT) or knock-out (KO). SKMel28R, have been obtained from SKMel28S by chronic exposure to non-lethal doses of BRAFi ^16^. R for BRAFi-resistant SKMel28 cells and S for sensitive. n**=**2 biologically independent experiments. Each histogram represents the mean ± s.d..

Together, our results support the hypothesis that AhR activity drives *SMAD3* expression along with the acquisition of BRAFi resistance.

### SMAD3 Drives Phenotype Switching and Resistance to Melanoma Therapies

To explore the mechanism underlying therapy sensitization upon SMAD3 inhibition, we examined the transcriptional program regulated by SMAD3 (Fig. 7). We hypothesized that the transcription factor SMAD3 may regulate the expression levels of several resistant genes and thereby induce a multifactorial effect, in which multiple drug resistance pathways are activated. We compared the SMAD3 ChIP-Seq ^45^ with those from BRAFi-resistance gene set established from three different sources, namely the enclosed screen, the screen performed in A375 ^8^, and other established BRAFi-resistance genes such as *AXL* or *NRP1* (Fig. 7A). A list of SMAD3-regulated genes was thus deduced, which comprises *SLIT2*, *RUNX2*, *NRP1*, *MMP2*, *JUNB*, *ITGB5*, *AXL*, *AFAP1* and *EGFR* (SMAD3-signature, Table S8). *SMAD3* depletion further validated *MMP2*, *AXL*, *EGFR* and *JUNB* as SMAD3-regulated genes in two melanoma cell lines (Fig. 7B). As expected, the SMAD3 activation by TGFβ exposure induced the SMAD3-signature in four melanoma cell lines (differentiated M229 & M249 *vs* dedifferentiated M238 & M238R) (Fig. 7C and D). Interestingly, the SMAD3-signature inducibility is higher in differentiated cells. Altogether, our results confirm that a SMAD3-regulated gene program is associated to therapy-resistance.

**Figure 7.**
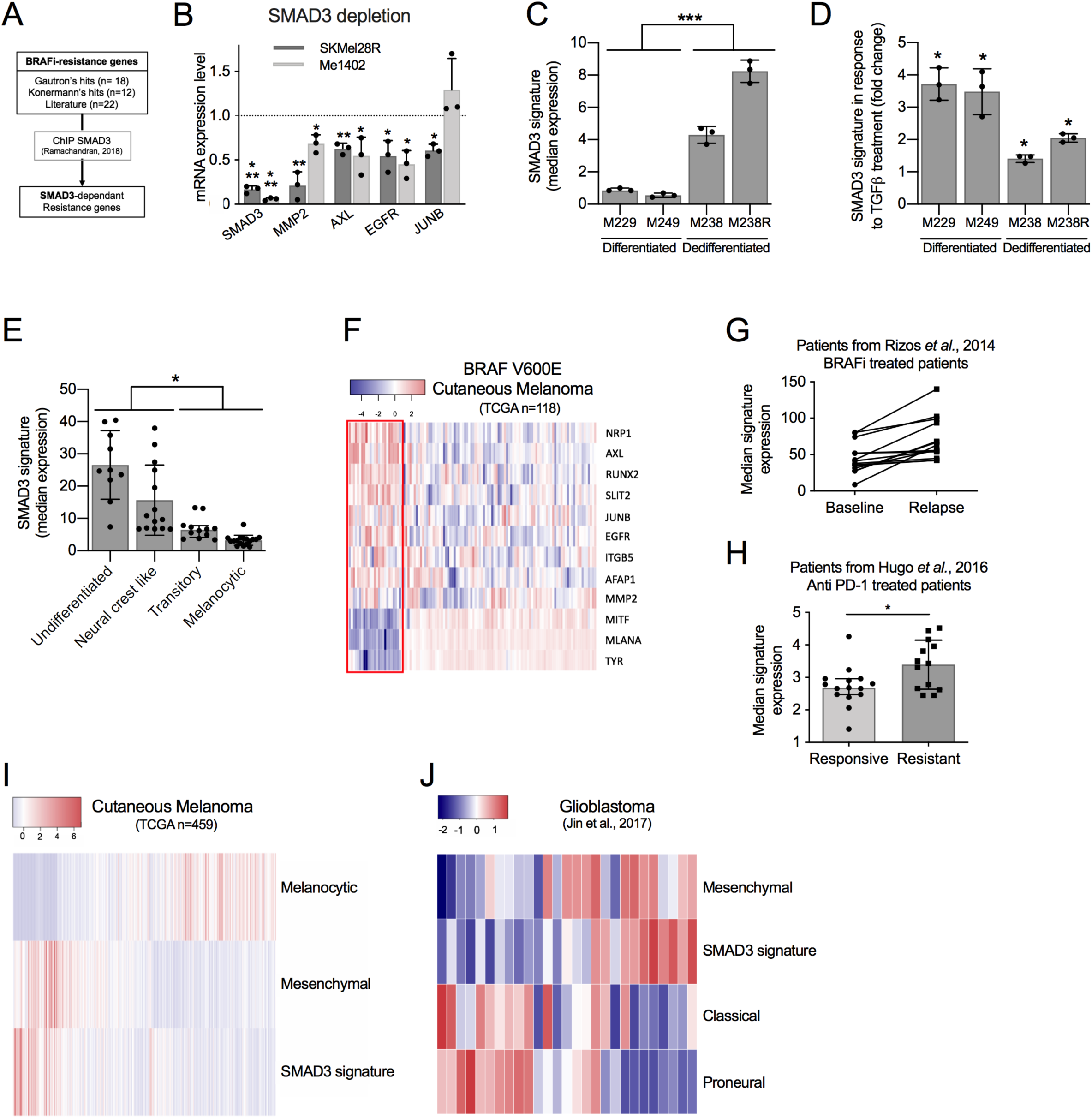
SMAD3 Drives Phenotype Switching and Resistance to Melanoma Therapies. (A) SMAD3 signature has been established by comparing BRAFi-resistance genes and SMAD3-regulated genes identified by chromatin immunoprecipitation followed by DNA sequencing (ChIP-Seq) ^45^. (B) SMAD3 depletion decreases expression of SMAD3-regulated genes. SMAD3 knock-down by siRNA decreased mRNA expression of BRAFi-resistance genes in SKMel28R and Me1402. n**=**3 biologically independent experiments. Each histogram represents the mean ± s.d. ; Bilateral Student test (with non-equivalent variances) *: p**<**0.05, **: p**<**0.01, ***: p**<**0.001. (C) Differentiation status determines basal expression level of SMAD3-signature. Basal SMAD3-signature in four melanoma cell lines grouped in function of their differentiation states ^17^. n**=**3 biologically independent experiments. Each histogram represents the median ± s.d. ; Bilateral Student test (with non-equivalent variances); ***: p<0.001 (D) Inducibility of SMAD3-signature in four melanoma cell lines exposed to TGF-β (10 ng/mL, 48h). Data were normalized to cell lines exposed to solvent (4mM HCl + 1mg/mL human BSA). n**=**3 biologically independent experiments. Each histogram represents the median ± s.d. ; Bilateral Student test (with non-equivalent variances); *: p<0.05 (E) SMAD3-signature discriminates differentiation states of 53 melanoma cell lines. The SMAD3-signature in four subgroups of melanoma cell lines ^17^. Each point represents a cell line; Each histogram represents the median with range ; Bilateral Student test (with non-equivalent variances); *: p<0.05 (F) Heatmap depicting mRNA levels of SMAD3-signature in BRAF(V600E) non-treated melanoma patients dataset from TCGA (SKCM, BRAF(V600E) mutated: n=118). Three pigmentation genes (*MITF*, *MLANA* and *TYR*) have been added to highlight the differentiation states of tumors. ∼20% of tumors are considered as dedifferentiated tumors (red box) with a high SMAD3-signature. (G) SMAD3-signature in two groups (before BRAFi treatment or during relapse) of V600E patients from the Rizos’s cohort ^66^. (H) SMAD3-signature in two groups (responsive to PD-1 or resistant) of patients with cutaneous melanoma exposed to PD-1 therapy (from the Hugo’s cohort) ^67^. Each histogram represents the median with range. Bilateral Student test (with non-equivalent variances) **: p**<**0.01. (I) SMAD3 signature overlapped with mesenchymal signature in melanoma tumors (TCGA, n=459) ^68^. Melanocytic signature highlights the differentiation states of tumors ^18^. SMAD3 signature and mesenchymal signature correlate in melanoma tumors (TCGA, n=459) ^47^. Pearson correlation test: p<0,0001. (J) The SMAD3 signature overlaps with the glioblastoma mesenchymal subtype ^48^. See also Fig. S4

We examined the SMAD3-signature in melanoma cell lines (n=53) (Fig. 7E) and cutaneous melanoma (n=118, TCGA cohort, tumors not exposed to targeted therapy) (Fig. 7F). The SMAD3-signature correlated with a dedifferentiation status as suggested above (Fig. 2H and 5B). Importantly, a subset of BRAF(V600E) melanoma patients (∼20%) expressing the SMAD3-signature was identified (Fig. 7F), indicating that these tumors contained dedifferentiated melanoma cells with potential intrinsic resistance to BRAFi. Therefore, these results indicated that the SMAD3-signature may be useful to identify a population of pre-existing BRAFi-resistant cells within drug-naive lesions.

To further illustrate the clinical relevance of our results, we assessed the expression levels of the SMAD3-signature in drug naive and drug resistant patients using publicly available dataset. We observed an increase in this SMAD3 signature in two-third of the patients exposed to BRAFi (Fig. 7G). Moreover, higher expression of the signature was also prominent in tumors that displayed resistance to immune checkpoint inhibitor therapy (PD-1) (Fig. 7H). Altogether, these results indicated that the SMAD3-signature is associated to resistance to both current melanoma therapies.

As the shift toward the mesenchymal-like state confers broad resistance to therapies ^46^, we postulated that the SMAD3-signature may be associated with this particular dedifferentiated phenotype. Comparing the SMAD3-signature with the classical mesenchymal-like signature ^47^ of melanoma (TCGA cohort) highlighted a significant correlation (Fig. 7I). To further confirm the link between the SMAD3-signature and a mesenchymal state, we searched for similarities with other mesenchymal states identified in two other cancers (glioblastoma [GBM] and hepatoma). As described for the cutaneous melanoma, different differentiation states have been characterized for GBM (proneural, classical, and mesenchymal GBM) ^48^. We found that the SMAD3-signature overlaps with the mesenchymal GBM signature (Fig. 7J). Mesenchymal GBM are the most aggressive GBM usually associated with poor overall survival ^4, 48^. The SMAD3-signature was also associated with epithelial-mesenchymal transition (EMT) in hepatoma (Fig. S4).

Altogether, these results indicate that the transcription factor SMAD3 and its downstream target genes confer resistance to targeted therapies by promoting a mesenchymal-like phenotype. Our work identifies AhR-SMAD3 axis as a target to overcome therapy resistance of melanoma (Fig. S5).

## DISCUSSION

Targeted therapy and immunotherapy have greatly improved the prognosis of patients with cancer, but resistance to these treatments restricts the overall survival of patients. Increasing evidence indicates that transcriptomic reprogramming is associated to persister cells emergence ^2, 3^ but the mechanism underlying resistance from this pool of cells remains elusive. Targeting tumor-promoting genes leading reprogramming could therefore constitute an attractive approach to prevent relapse, at least in some specific contexts ^1^. Here, using a whole genome approach, we searched for pathways that trigger the transcriptional reprogramming of persister cells into drug-resistant cells. Based on CRISPR screen, data mining and *in vivo* experiments, we identified and validated three genes (*SMAD3*, *BIRC3* and *SLC9A5*) able to promote both BRAFi-resistance and tumor growth. Our work expands our understanding of the biology of persister cells and highlight novel drug vulnerabilities that can be exploited to develop long-lasting antimelanoma therapies.

Even if CRISPR-SAM screen is a leading-edge genetic tool, several concerns must be considered. As observed for all screening approaches, false positives and false negatives are engendered rendering the validation experiments a crucial step. In this study, we clearly showed that the number of sgRNAs per target is an important parameter. For our best hit, *EGFR*, only two sgRNAs were enriched in BRAFi-treated cells. Thus, it is highly likely that we missed interesting BRAFi-resistance genes (false negatives) due to the number of sgRNA/gene (at least 3 sgRNAs/gene in this library). A recent publication confirmed that sgRNAs are not all functional in CRISPRa libraries and it could be interesting to increase the number of sgRNAs per target and to cover more TSS per gene ^33^. Interestingly, the sgRNA library used in our study displays for several genes up to 27 sgRNAs. These sgRNAs target different isoforms (or TSS) of these genes. By examining the 12 sgRNAs targeting *SMAD3*, we found that only 2 sgRNA are enriched. These two sgRNAs promote the expression of the longest *SMAD3* isoform; the only *SMAD3* mRNA expressed in our model (501Mel cells) (data not shown). Based on these observations, we believe that mRNA isoforms identified by RNA-sequencing should be considered during the sgRNAs selection for each model. By this way, it would be easy to disqualify a part of sgRNAs (without transactivating effect).

The second lesson of this CRISPR screen is the weak sgRNAs enrichment in BRAFi-exposed cells. Except for *EGFR*, the sgRNAs enrichment is about 2. Despite these values, we confirmed that these candidates seem robust such as SMAD3 (*in vitro*, *in vivo* and in patients). It is tempting to explain this fact by the duration of treatment (BRAFi exposure) and the dose (2μM). By increasing the dose (i.e. 5μM), we showed that BRAFi killed more than 95% of 501Mel cells in four days (Fig. 5M). So, this protocol is not achievable because it would induce too many false negatives. The other option consists to increase the duration of BRAFi treatment (and keep a low BRAFi concentration i.e. 2μM). However, a recent publication demonstrated that a long-term BRAFi exposure promotes a dedifferentiation process conferring BRAFi resistance^3, 49^. So, this alternative protocol could be perilous by inducing cell resistance to BRAFi independently of the sgRNA expression. Here, we selected a melanoma cell model exhibiting a differentiated profile as the vast majority of metastatic melanoma tumors (89% in the TCGA cohort), a short period of treatment (14 days) and an intermediate dose of BRAFi (2μM). In fact, we followed, except the cell line, the protocol established by Feng Zhang’s lab, who developed the CRISPR-SAM library ^8^. The differentiation status of the cell line could be important since the transactivation mediated by CRISPR-SAM is possible only for active promoter in basal condition and the magnitude of transactivation relies on the basal expression level. Here, we showed that 501Mel cells express low level of SMAD3 in basal condition and the transactivation obtained by CRISPR-SAM is massive (Fig. 1I). Moreover, SMAD3 is a transcription factor which promotes the expression of various genes including potent BRAFi-resistance genes (AXL and EGFR). Thus, a robust transactivation of genes encoding a transcription factor, a transporter or an enzyme is more inclined to be enriched with our protocol, especially if the basal gene expression is low. Here, we identified the transcription factor SMAD3 and the transporter SLC9A5 (Sodium/hydrogen exchanger 5), validated *in vitro* and *in vivo*.

BRAFi-resistance relies, at least in part, on the phenotypic plasticity of melanoma cells ^6, 17, 32^. These cells may escape the deleterious effect of drug combinations such as BRAFi + MEKi. Among these cells, those harboring a mesenchymal-like phenotype (usually named invasive cells) display high intrinsic resistance to MAPK therapeutics ^22, 23, 25, 32^. Enrichment in AXL^high^ subpopulation (considered as invasive and mesenchymal-like cells) is a common feature of drug-resistant melanomas ^23, 25^. Targeting mesenchymal-like cells using an antibody-drug conjugate, AXL-107-MMAE, showed promising effects in a preclinical model of melanoma ^24^. The emergence of AXL^high^ cells is currently explained by the decrease in *MITF* activity, but the mechanism of resistance to MAPK therapeutics remains unclear. Here, we demonstrate that the AhR-SMAD3 axis governs the expression levels of potent BRAFi-resistance genes, including *AXL*, *EGFR*, and *MMP2*.

The dedifferentiation process, conferring BRAFi-resistance, requires transcriptomic reprogramming by transcription factors. Retinoic acid receptor gamma (RXRγ) was identified as a crucial transcription factor that promotes the emergence of the drug-tolerant subpopulation of NCSCs ^6^. An increase in AXL^high^-positive cell population was reported following MAPK inhibition in the presence of an RXRγ antagonist. This increase may explain why this co-treatment only delays but does not completely prevent relapse in PDXs. This confirms the need to develop strategies that prevent melanoma dedifferentiation during BRAFi treatment. Thus, our data indicating that SMAD3 is a key transcriptional factor involved in the emergence of drug-resistant mesenchymal-like cells in response to MAPK remains of great interest. Other transcription factors have been associated to BRAFi resistance such as JUN ^50^ and AhR ^18, 51^. We recently showed that a sustained AhR activation promotes the dedifferentiation of melanoma cells and the expression of BRAFi-resistance genes ^18^. As proof-of concept, we demonstrated that differentiated and BRAFi-sensitive cells can be directed towards an AhR-dependent resistant program using AhR agonists. To abrogate the deleterious AhR sustained-activation, we identified Resveratrol, a clinically compatible AhR-antagonist. Combined with BRAFi, Resveratrol reduces the number of BRAFi-resistant cells and delays relapse.

However, because its poor bioavailability, this AhR antagonist ^52^ is not curative ^18^. New AhR antagonists are currently being evaluated (Hercules Pharmaceuticals and Ikena Oncology). Another option overcoming this pharmacological caveat would be to target actors, downstream of the AhR pathway. Consistently with these results, here, we demonstrated that AhR drives the expression level of *SMAD3* and the SMAD3-regulated gene program promotes therapy-resistance in cutaneous melanoma. Thus, we propose an AhR-SMAD3 impairment as a strategy to overcome melanoma resistance. Recently, conditional deletion of Smad7, a negative regulator of TGF-β/SMAD pathway, led to sustained melanoma growth and at the same time promoted massive metastasis formation ^53^, confirming that TGF-β/SMAD pathway is a promising target for melanoma ^54^. In addition, Rizos’s team further illustrated the link between the TGF-β and melanoma therapy resistance. They showed that TGF-β promotes a de-differentiation phenotype, which is a common mechanism of resistance to PD-1 inhibitors ^55^.

Several questions remain unsolved. We previously reported that activated AhR reprograms the transcriptome of melanoma cells mediating BRAFi-resistance. In this study, we demonstrate that a SMAD3-regulated gene expression program promotes therapy-resistance in cutaneous melanoma and EMT. Importantly, *SMAD3* expression levels during resistance acquisition is dependent, at least in part, on AhR. Thus, it would be interesting to precisely define the role of AhR and SMAD3 in the induction of each BRAFi-resistance gene. To date, no physical interaction between AhR and SMAD3 proteins has been reported, suggesting that AhR acts as an upstream regulator of SMAD3 axis. It is noteworthy that the increased expression levels of SMAD3 by AhR expands the possibility of fine tuning gene expression since SMAD3 interacts with SMAD2 but also with JUN, TEADs and YAP1 ^56–58^. Because these three transcription factors have also been associated to therapies-resistance in melanoma ^16, 22, 59, 60^, our results suggest that regulation of BRAFi-resistance genes expression is multiparametric and probably more sophisticated than initially though. Nonetheless, the elucidation of these transcriptional programs and networks governing BRAFi-resistance genes and relapse is important for optimal target selection and the development of rationale and effective combination strategies.

BRAFi resistance may be achieved through the exposure of melanoma cells to TGF-β, demonstrating that transcriptome reprogramming may confer resistance without the need for pre-existing or *de novo* mutations ^61^. The TGF-β pathway promotes a shift toward the mesenchymal state ^62^. The resulting dedifferentiation modifies the expression of the adhesion molecules in the cell, supporting a migratory and invasive behavior. Together, our results strongly indicate that the SMAD3-regulated genes are critical players in melanoma resistance to therapies by promoting an EMT-like process. EMT reversal represents a powerful approach, as it may reduce the invasive behavior of cancer cells and favor re-differentiation, synonymous of a decrease in BRAFi-resistance gene expression ^63, 64^. By combining anti-EMT drug and targeted therapy such as SMAD3i and BRAFi, we efficiently reduced amount of persister cells. We anticipate that SMAD3 inhibition should limit the risk of resistance to therapies, since a decrease of expression levels of several BRAFi-resistance genes is obtained with SMAD3i (SIS3). SMAD3 inhibition is expected to be more efficient than inhibitors targeting single downstream targets such as AXL or EGFR. In conclusion, our work highlights novel drug vulnerabilities that can be exploited to develop long-lasting antimelanoma therapies.

Given the plasticity of melanoma cells and the capability of tumor microenvironment to produce TGF-β ^65^, our work also warrant further investigation of the source of TGF-β as another approach to prevent acquisition of the therapy-resistant mesenchymal phenotype.

## ACKNOWLEDGEMENTS

The authors thank the Gene Expression and Oncogenesis team from the CNRS UMR6290 especially M. Migault, A. Forestier, and L. Boussemart; the Rennes FHU CAMIn team; and E. Watrin and C. Hitte for providing scientific expertise.

The authors acknowledge the SFR Biosit core facilities of Rennes 1 University with the ARCHE animal housing facility, the cell culture L3 facility and the Human and Environmental Genomics platform for their help and support.

This study received financial support from the following: Fondation ARC pour la Recherche; Ligue Nationale Contre le Cancer (LNCC); Départements du Grand-Ouest; Région Bretagne; University of Rennes 1; CNRS; SFR Biosit, Association Vaincre le Cancer.

Further support was provided by a “Ligue Nationale Contre le Cancer” (LNCC) Grand Ouest fellowship (AG) and from the Région Bretagne (AG), and Fondation ARC pour la Recherche (AG) and the LNCC (AQ) and from French Ministry of Research (NT) and from Institut National contre le Cancer (INCa) (AP). The authors are grateful to Feng Zhang for providing the Human CRISPR 3-plasmid activation pooled library (SAM) (Addgene).

## AUTHOR CONTRIBUTIONS

Conceptualization: AG & DG.

Methodology: AG, LB, AQ, MA & DG.

Software: AG, MA, SC, CC & FR.

Formal analysis: AG, LB, AQ, MA, SC, CC & FR.

Investigation: AG, LB, AQ, AP, MA, SC, AP, HML, NT & DG.

Writing-original draft: AG & DG.

Writing-Review & Editing: AG, DG, LB, AQ, SC, MDG, FR, MA, CC & JCM.

Visualization: AG, LB, AQ, MA, SC, FR, CC & DG.

Supervision: DG.

Project Administration: DG & MDG.

Funding: DG, SC & MDG.

## DATA AVAILABILITY

The datasets generated during and/or analysed during the current study are available from the corresponding author on reasonable request. CRISPRa screen data have been uploaded to the ArrayExpress (https://www.ebi.ac.uk/arrayexpress/) under accession code E-MTAB-8595. Statistics source data are available in Table S10 and unprocessed original blots are shown in Fig. S6

## METHODS

### Reagents

- DMSO - Sigma-Aldrich (D8418)
- BRAF inhibitors: Vemurafenib (PLX4032) - Selleckchem (RG7204); Paradox Breaker (PLX8394) - MedChem Express (HY-18972)
- SMAD3 inhibitor: SIS3 - SantaCruz Biotechnology (sc-222318)
- 2,3,7,8-TCDD (TCDD) - Sigma-Aldrich, (48599)
- 2-(1*H*-Indol-3-ylcarbonyl)-4-thiazolecarboxylic acid methyl ester (ITE) - MedChemExpress (HY-19317)
- CH-223191 - Selleckchem (S7711)
- TGF-β recombinant - SantaCruz Biotechnology (240-B-010)

### Cell Lines and Culture Conditions

501Mel, Me1402 and HEK293T cell lines were obtained from ATCC. SKMel28 S & R cell lines were obtained from J.C Marine’s lab at VIB Center for Cancer Biology, VIB, Leuven, Belgium. 501Mel and SKMel28 AhR knock-out cell lines have been established as previously described ^18^. M229S, M229R, M238S, M238R & M249 were obtained from Thomas Graeber’s lab at department of Molecular and Medical Pharmacology, University of California, Los Angeles, USA. All melanoma cell lines were grown in humidified air (37°C, 5% CO2) in RPMI-1640 medium (Gibco BRL, Invitrogen, Paisley, UK) supplemented with 10% Fetal Bovine Serum (FBS) (PAA cell culture company) and 1% Penicillin-Streptomycin (PS) antibiotics (Gibco, Invitrogen). HEK293T were grown in DMEM (Gibco BRL, Invitrogen, Paisley, UK) supplemented as melanoma cell lines media. SKMel28R, M229R and M238R are cultivated in presence of 1μM vemurafenib. All cell lines have been routinely tested for mycoplasma contamination (Mycoplasma contamination detection kit; rep-pt1; InvivoGen - San Diego - CA).

### CRISPR-SAM screens

A detailed protocol is available as supplementary file. Briefly, lentiviral productions have been performed as recommended (http://tronolab.epfl.ch), using HEK293T cells, psPAX2, pVSVG and vectors required for CRISPR-SAM according to Zhang lab ^69^. Infections were performed overnight in presence of 4 or 8μg of polybrene per ml. All vectors have been provided by Addgene. SgRNA Library was amplified and prepared as described by Zhang Lab ^69^. 501Mel cells were transduced to stably express dCAS-VP64 (cat. no. 61425) and MS2-P65-HSF1 (cat. no. 61426), before to transduce them with SAM sgRNA library (lentiSAMv2, 3-plasmid system, cat. no. 1000000057) at a MOI of 0.2. Infected cells have been selected using antibiotics: Blasticidin (2 μg/mL, 5 days), Hygromycin B (200 μg/mL, 5 days), Zeocine (600 μg/mL, 5 days). *In vitro* CRISPR-SAM screens (Fig. 2) were conducted as described by Zhang lab ^69^. The 501Mel cells expressing the sgRNA library were split into three groups: DMSO (solvent for BRAFi), vemurafenib (PLX4032, 2μM, Selleckchem), Paradox Breaker (PLX8394, 2μM, MedChemExpress). After 14 days, resistant cell populations have been amplified. A minimum of 36×10^6^ cells has been pellet and further sgRNA enrichment analysis. Cell libraries have been cryopreserved at −80°C for further *in vivo* experiments. The results presented are pooled data from two independent screens. To generate 501Mel cells overexpressing individually *SMAD3*, *BIRC3* or *SLC9A5*, 501Mel expressing dCAS-VP64 and MS2-P65-HSF1 were transduced to stably express specific sgRNAs (Table S9). Infected cells have been selected using zeocin (600 μg/mL, 5 days). Manipulations of lentivirus were performed in the biosafety level 3 containment laboratory core facility of the Biology and Health Federative Research Structure of Rennes (Biosit)

### Xenograft

Mice were maintained under specific pathogen-free conditions in our accredited animal house (A 35_238_40). The animal study follows the 3R (replace_reduce_refine) framework and has been filed with and approved by the French Government Board (No. 04386.03). Animal welfare is a constant priority: animals were thus euthanized under anaesthesia.

To identify tumor-promoting genes *in vivo* (Fig. 1D), two cell populations were subcutaneously xenografted on female NMRI nude mice flanks (3×10^6^ cells per mouse); 501Mel (6 mice) and 501Mel CRISPR-SAM cell library (10 mice). Tumor growth was assessed as previously described ^70^, during 20 weeks. After mice sacrifice, tumors were sampled and conserved at −80°C for further sgRNA enrichment analysis.

For individual validation of tumor-promoting genes (Fig. 1H-K): 3×10^6^ of 501Mel cells overexpressing either *SMAD3*, *BIRC3* or *SLC9A5* were subcutaneously xenografted on female NMRI nude mice flanks (6 mice per group). In each group, mice were injected on both flanks: sgRNA 1 on right flank and sgRNA 2 on left flank (see Table S9 for sgRNA sequences). Tumor growth was assessed as previously described ^70^ until the endpoint (600mm^3^).

In order to identify BRAFi-resistance and tumor growth genes *in vivo* (Fig. 4), three cell populations were subcutaneously xenografted on female NMRI nude mice flanks (3×10^6^ cells per mice); 501Mel (6 mice), 501Mel CRISPR-SAM vemurafenib resistant (10 mice), 501Mel CRISPR-SAM Paradox Breaker resistant (12 mice). Tumor growth was assessed as previously described ^70^, during 20 weeks. After mice sacrifice, tumors were sampled and conserved at −80°C for further sgRNA enrichment analysis.

These experiments are compliant with all relevant ethical regulations regarding animal research.

### sgRNA enrichment analysis

Genomic DNAs from cell pellets (>36.10^6^ cells) and tumors (>400mg) were extracted using the Zymo Research Quick-gDNA MidiPrep according to the manufacturer’s protocol. PCR amplifications and quality controls have been done as described by Zhang lab ^69^.

### sgRNA Sequencing

Sequencing was performed by the Human & Environmental Genomics core facility of Rennes on a HiSeq 1500 (Rapid SBS kit v2 1×100 cycles, Illumina). Base Calling was performed with Illumina’s CASAVA pipeline (Version 1.8).

### Bioinformatic Analysis of sgRNA and Gene Hits

Data processing was conducted using the MAGeCK v0.5.6 software ^71^. Briefly, read counts from different samples are first median-normalized to adjust for the effect of library sizes and read count distributions (mageck count with option: --norm-method median). Then, in an approach similar to those used for differential RNA-Seq analysis, the variance of read counts is estimated by sharing information across features and a negative binomial model is used to test whether sgRNA abundance differs significantly between the treatment conditions and the DMSO control. Positively or negatively selected sgRNA are ranked according to adjusted P-values (false discovery rate) and gene log fold changes computed with the modified robust ranking aggregation algorithm implemented in MAGeCK (mageck test with options: --norm-method median, --gene-lfc-method alphamedian, --adjust-method fdr).

### RNA interference

All siRNAs were transfected at 66nM using Lipofectamine RNAiMAX (Invitrogen). For survival assay, 4.000 cells were seeded in 96-well plates, in quadruplicates. The following day, cells have been FBS-starved (1% FBS overnight). Thirty-six hours after transfection, cells were exposed to DMSO or BRAFi (vemurafenib (PLX4032); Selleckchem; 1μM). After 84h of treatment, cell density was measured by methylene blue assay, as previously described ^72^. For RNA analysis, 50.000 cells were seeded in 12-well plates. Cells were harvested 48h after transfection. All siRNAs were purchased from IDT DNA (Table S9).

### Treatment experiments

8,000 (SKMel28R) or 12,000 (Me1402) cells were seeded in 96-well plates, in quadruplicates. For SMAD3i + vemurafenib combination treatment: 5,000 (501Mel) or 8,000 (M249, SKMel28R) cells were seeded in 96-well plates, in triplicates. Cells were exposed, 6h after seeding, to either DMSO, BRAFi (vemurafenib; 5μM) or the combination BRAFi (5μM) + SMAD3i (0, 3, 10, 15 or 20μM or solvent) for 4 days. Cell density was evaluated using methylene blue assay, as previously described ^72^.

### Half maximal inhibitory concentrations (IC_50_ values)

Cell sensitivity to vemurafenib or SMAD3i (SIS3) has been established by cell density measurement and calculation of the IC_50_ using GraphPad (PRISM8.0®) as previously described^18^. 5,000 (501Mel) or 8,000 cells (other cell lines) were plated and exposed to inhibitor(s) at indicated concentrations for 4 days. Cells have been exposed to inhibitors 6 hours after plating.

### Melanoma spheroids

Spheroids were prepared using the liquid overlay method. Briefly, 500 μL of melanoma cells (10,000 cell/mL) were added to a 24-well plate coated with 1.5% agar (Invitrogen). Plates were left to incubate for 72 hours; by this time, cells had organized into 3-dimensional (3D) spheroids. Spheroids were then harvested and added into 1 ml of a solution of collagen I (2 mg/ml - Corning) with MEM 1X (Gibco), acetic acid 0,02N and neutralization buffer (HEPES 200mM pH 7.4; sodium bicarbonate 2.2%; NaOH 0.2N). The suspension was then added to a 12-well plate coated with 1.5% agar. Normal medium was overlaid on top of the solidified collagen after 2 hours of incubation. After 48 hours, medium was renewed. Pictures of the invading spheroids were monitored at different times using a Zeiss microscope.

### RNA extraction & RT-qPCR expression

Experiments have been done as previously described ^70^. Primers used for RT-qPCR experiments are available in Table S10.

### Western-blot experiments

Experiments were performed as previously described ^70^. Membranes were probed with suitable antibodies and signals were detected using the LAS-3000 Imager (Fuji Photo Film). The primary and secondary antibodies are described in Table S9.

### SMAD-luciferase Assay

Experiments are based on the SMAD responsive element: Cignal Lenti SMAD Reporter (luc) (CLS-017L from Quiagen) and as control the Cignal Lenti Negative Control (luc). The SMAD-responsive luciferase vector encodes the Firefly luciferase reporter gene under the control of a minimal (m)CMV promoter and tandem repeats of the SMAD Binding Element (SBE). Melanoma cells line were infected as previously described ^70^ and infected cells were selected using antibiotic selection (puromycin 2μg/mL, 7 days). Cells were exposed to TGFβ +/- SMAD3i as detailed in Figure 5K legend. Luciferase assays were then performed with a Promega kit according to the manufacturer’s instructions. Data were expressed in arbitrary units, relative to the value of luciferase activity levels found in TGFβ-exposed cells, arbitrarily set at 100%. Firefly luciferase activity was normalized to protein content using Bicinchoninic Acid Kit from Sigma-Aldrich® and measured with using a luminometer (Centro XS3 LB960, Berthold Technologies).

### *In silico* analyses

Heatmaps were generated with R-packages heatmap3 ^73^.

Gene Set Enrichment Analysis were performed using the Broad Institute software.

SKCM TCGA expression data were obtained using OncoLNC portal (http://www.oncolnc.org; ^74^). Cell state categorization into four differentiation states (Undifferentiated, Neural crest-like, Transitory, Melanocytic) of SKCM TCGA tumors and were performed using expression data of previously established gene sets ^17^.

Genomic alterations of SKCM TCGA tumors were analyzed using cBioPortal (http://www.cbioportal.org).

CCLE cell lines expression data and IC_50_ were obtained from GSE36139 and from the original publication ^34^, respectively.

Expression data of 501Mel were obtained from our previously published RNAseq (GSE95589)_70_.

Expression data of M229, M238 and SKMel28 at different resistance steps were obtained from GSE75313 ^13^.

Patients median expression of our 18 hits in baseline *vs* relapse (BRAFi) are based on expression data from GSE65186 ^16^ and GSE50509 ^66^.

Expression data of invasive *vs* proliferative cell lines were obtained from GSE60666 ^22^. Analysis of our 18 hits expression in EGFR-positive sorted cells was realized based on GSE97682 ^32^.

Analysis of *SMAD3*, *BIRC3* and *EGFR* expression among 52 cell lines previously categorized as Undifferentiated, Neural crest-like, Transitory or Melanocytic were performed using http://systems.crump.ucla.edu/dediff and GSE80829.

SMAD3 ChIP-Seq data have been obtained from GSE92443 ^45^.

The comparison of the median SMAD3 signature in anti-PD-1 responsive *vs* resistant patients has been performed *via* expression data from GSE78220 ^67^.

SMAD3 signature median expression was compared to melanocytic one (*MITF*, *OCA2*, *MLANA*, *TYR*, *DCT*) and mesenchymal one (52 genes; ^47^) *via* expression data from SKCM TCGA obtained from OncoLNC.

The comparison between median expression of SMAD3 signature with classical, proneural and mesenchymal ones in a cohort of glioblastoma patients was based on expression data from GSE103366 ^48^.

Gene Set Enrichment Analysis on Huh28 +/- TGF-β was realized based on GSE102109 ^75^. Gene Set Enrichment Analysis on 3sp cells (mesenchymal) *versus* 3p cells (epithelial) was realized based on GSE26391 ^76^.

Single cell RNAseq data of a BRAF mutant PDX model undergoing BRAF and MEK inhibition were retrieved (GSE116237). The 674 single cells were projected into a two-dimensional space using t-distributed stochastic neighbor embedding (tSNE) and colored according to their drug tolerant state (DTC) identity ^6^. In a second step, the activity of the resistance gene expression signature (18 genes) was quantified per cell using the AUCell algorithm ^77^ resulting into an AUCell score (0<range<0.4), which was used to gradient-color the tSNE plot. Finally, the activity of the resistance gene signature was quantified per DTC population using Graphpad (****p<0.0001, Mann-Whitney test).

### Statistical analyses

Data are presented as mean ± s.d. unless otherwise specified, and differences were considered significant at a p value of less than 0.05. Comparisons were performed using Bilateral Student test (with non-equivalent variances), One-way ANOVA and Sidak’s multiple comparisons test or Pearson Correlation as specified in figure legends. All statistical analyses were performed using Prism 8 software (GraphPad, La Jolla, CA, USA) or Microsoft Excel software.

## REFERENCES

1. Bailey, M. H. et al. Comprehensive Characterization of Cancer Driver Genes and Mutations. Cell 174, 1034–1035 (2018).

2. Puisieux, A., Brabletz, T. & Caramel, J. Oncogenic roles of EMT-inducing transcription factors. Nat. Cell Biol. 16, 488–94 (2014).

3. Bai, X., Fisher, D. E. & Flaherty, K. T. Cell-state dynamics and therapeutic resistance in melanoma from the perspective of MITF and IFNγ pathways. Nat. Rev. Clin. Oncol. (2019). doi:10.1038/s41571-019-0204-6

4. Patel, A. P. et al. Single-cell RNA-seq highlights intratumoral heterogeneity in primary glioblastoma. Science 344, 1396–1401 (2014).

5. Tirosh, I. et al. Dissecting the multicellular ecosystem of metastatic melanoma by single-cell RNA-seq. Science (80-.). 352, 189–196 (2016).

6. Rambow, F. et al. Toward Minimal Residual Disease-Directed Therapy in Melanoma. Cell 174, 843–855.e19 (2018).

7. Shalem, O., Sanjana, N. E. & Zhang, F. High-throughput functional genomics using CRISPR–Cas9. Nat. Rev. Genet. 16, 299–311 (2015).

8. Konermann, S. et al. Genome-scale transcriptional activation by an engineered CRISPR-Cas9 complex. Nature 517, 583–588 (2015).

9. Meyers, R. M. et al. Computational correction of copy number effect improves specificity of CRISPR-Cas9 essentiality screens in cancer cells. Nat. Genet. 49, 1779–1784 (2017).

10. Behan, F. M. et al. Prioritization of cancer therapeutic targets using CRISPR-Cas9 screens. Nature (2019). doi:10.1038/s41586-019-1103-9

11. Ascierto, P. A. et al. Cobimetinib combined with vemurafenib in advanced BRAFV600-mutant melanoma (coBRIM): updated efficacy results from a randomised, double-blind, phase 3 trial. Lancet Oncol. 17, 1248–1260 (2016).

12. Welsh, S. J., Rizos, H., Scolyer, R. A. & Long, G. V. Resistance to combination BRAF and MEK inhibition in metastatic melanoma: Where to next? Eur. J. Cancer 62, 76–85 (2016).

13. Song, C. et al. Recurrent Tumor Cell-Intrinsic and -Extrinsic Alterations during MAPKi-Induced Melanoma Regression and Early Adaptation. Cancer Discov. 7, 1248–1265 (2017).

14. Sullivan, R. J. & Flaherty, K. T. Resistance to BRAF-targeted therapy in melanoma. Eur. J. Cancer 49, 1297–1304 (2013).

15. Wagle, N. et al. Dissecting therapeutic resistance to RAF inhibition in melanoma by tumor genomic profiling. J. Clin. Oncol. Off. J. Am. Soc. Clin. Oncol. 29, 3085–3096 (2011).

16. Hugo, W. et al. Non-genomic and Immune Evolution of Melanoma Acquiring MAPKi Resistance. Cell 162, 1271–1285 (2015).

17. Tsoi, J. et al. Multi-stage Differentiation Defines Melanoma Subtypes with Differential Vulnerability to Drug-Induced Iron-Dependent Oxidative Stress. Cancer Cell 0, (2018).

18. Corre, S. et al. Sustained activation of the Aryl hydrocarbon Receptor transcription factor promotes resistance to BRAF-inhibitors in melanoma. Nat. Commun. 9, 4775 (2018).

19. Talebi, A. et al. Sustained SREBP-1-dependent lipogenesis as a key mediator of resistance to BRAF-targeted therapy. Nat. Commun. 9, 2500 (2018).

20. Rapino, F. et al. Codon-specific translation reprogramming promotes resistance to targeted therapy. Nature 558, 605–609 (2018).

21. Hoek, K. S. & Goding, C. R. Cancer stem cells versus phenotype-switching in melanoma. Pigment Cell Melanoma Res. 23, 746–759 (2010).

22. Verfaillie, A. et al. Decoding the regulatory landscape of melanoma reveals TEADS as regulators of the invasive cell state. Nat. Commun. 6, 6683 (2015).

23. Konieczkowski, D. J. et al. A melanoma cell state distinction influences sensitivity to MAPK pathway inhibitors. Cancer Discov. 4, 816–827 (2014).

24. Boshuizen, J. et al. Cooperative targeting of melanoma heterogeneity with an AXL antibody-drug conjugate and BRAF/MEK inhibitors. Nat. Med. 24, 203–212 (2018).

25. Müller, J. et al. Low MITF/AXL ratio predicts early resistance to multiple targeted drugs in melanoma. Nat. Commun. 5, 5712 (2014).

26. Smith, M. P. et al. Inhibiting Drivers of Non-mutational Drug Tolerance Is a Salvage Strategy for Targeted Melanoma Therapy. Cancer Cell 29, 270–284 (2016).

27. Nassar, D. & Blanpain, C. Cancer Stem Cells: Basic Concepts and Therapeutic Implications. Annu. Rev. Pathol. Mech. Dis. 11, 47–76 (2016).

28. Halaban, R. et al. PLX4032, a selective BRAFV600Ekinase inhibitor, activates the ERK pathway and enhances cell migration and proliferation of BRAFWTmelanoma cells. Pigment Cell Melanoma Res. 23, 190–200 (2010).

29. Ohanna, M. et al. Senescent cells develop a PARP-1 and nuclear factor-B-associated secretome (PNAS). Genes Dev. 25, 1245–1261 (2011).

30. Lamar, J. M. et al. The Hippo pathway target, YAP, promotes metastasis through its TEAD-interaction domain. Proc. Natl. Acad. Sci. 109, E2441–E2450 (2012).

31. Sun, C. et al. Reversible and adaptive resistance to BRAF(V600E) inhibition in melanoma. Nature 508, 118–122 (2014).

32. Shaffer, S. M. et al. Rare cell variability and drug-induced reprogramming as a mode of cancer drug resistance. Nature 546, 431–435 (2017).

33. Sanson, K. R. et al. Optimized libraries for CRISPR-Cas9 genetic screens with multiple modalities. Nat. Commun. 9, 5416 (2018).

34. Barretina, J. et al. The Cancer Cell Line Encyclopedia enables predictive modeling of anticancer drug sensitivity. Nature 483, 603–607 (2012).

35. Rizzolio, S. et al. Neuropilin-1 upregulation elicits adaptive resistance to oncogene-targeted therapies. J. Clin. Invest. 128, 3976–3990 (2018).

36. Levy, C., Khaled, M. & Fisher, D. E. MITF: master regulator of melanocyte development and melanoma oncogene. Trends Mol. Med. 12, 406–414 (2006).

37. Moriceau, G. et al. Tunable-combinatorial mechanisms of acquired resistance limit the efficacy of BRAF/MEK cotargeting but result in melanoma drug addiction. Cancer Cell 27, 240–256 (2015).

38. Gao, J. et al. Integrative Analysis of Complex Cancer Genomics and Clinical Profiles Using the cBioPortal. Sci. Signal. 6, pl1–pl1 (2013).

39. Prahallad, A. et al. Unresponsiveness of colon cancer to BRAF(V600E) inhibition through feedback activation of EGFR. Nature 483, 100–3 (2012).

40. Berking, C. et al. Transforming Growth Factor-b1 Increases Survival of Human Melanoma through Stroma Remodeling. Cancer Res. 8306–8316 (2001).

41. Jinnin, M., Ihn, H. & Tamaki, K. Characterization of SIS3, a Novel Specific Inhibitor of Smad3, and Its Effect on Transforming Growth Factor-beta1-Induced Extracellular Matrix Expression. Mol. Pharmacol. 69, 597–607 (2006).

42. Chihara, Y. et al. A small-molecule inhibitor of SMAD3 attenuates resistance to anti-HER2 drugs in HER2-positive breast cancer cells. Breast Cancer Res. Treat. 166, 55–68 (2017).

43. Yang, S. Y., Ahmed, S., Satheesh, S. V & Matthews, J. Genome-wide mapping and analysis of aryl hydrocarbon receptor (AHR)- and aryl hydrocarbon receptor repressor (AHRR)-binding sites in human breast cancer cells. Arch. Toxicol. 92, 225–240 (2018).

44. Lo, R. & Matthews, J. High-resolution genome-wide mapping of AHR and ARNT binding sites by ChIP-Seq. Toxicol. Sci. 130, 349–61 (2012).

45. Ramachandran, A. et al. TGF-β uses a novel mode of receptor activation to phosphorylate SMAD1/5 and induce epithelial-to-mesenchymal transition. Elife 7, 1–29 (2018).

46. Redfern, A. D., Spalding, L. J. & Thompson, E. W. The Kraken Wakes: induced EMT as a driver of tumour aggression and poor outcome. Clin. Exp. Metastasis 35, 285–308 (2018).

47. Mak, M. P. et al. A Patient-Derived, Pan-Cancer EMT Signature Identifies Global Molecular Alterations and Immune Target Enrichment Following Epithelial-to-Mesenchymal Transition. Clin. Cancer Res. An Off. J. Am. Assoc. Cancer Res. 22, 609–620 (2016).

48. Jin, X. et al. Targeting glioma stem cells through combined BMI1 and EZH2 inhibition. Nat. Med. 23, 1352–1361 (2017).

49. Tsoi, J. et al. Multi-stage Differentiation Defines Melanoma Subtypes with Differential Vulnerability to Drug-Induced Iron-Dependent Oxidative Stress. Cancer Cell 1–15 (2018).

50. Titz, B. et al. JUN dependency in distinct early and late BRAF inhibition adaptation states of melanoma. Cell Discov. 2, 16028 (2016).

51. Liu, Y. et al. Blockade of IDO-kynurenine-AhR metabolic circuitry abrogates IFN-γ-induced immunologic dormancy of tumor-repopulating cells. Nat. Commun. 8, 15207 (2017).

52. Santos, A. C., Veiga, F. & Ribeiro, A. J. New delivery systems to improve the bioavailability of resveratrol. Expert Opin. Drug Deliv. 8, 973–990 (2011).

53. Tuncer, E. et al. SMAD signaling promotes melanoma metastasis independently of phenotype switching. J. Clin. Invest. (2019). doi:10.1172/JCI94295

54. Javelaud, D. et al. Stable overexpression of Smad7 in human melanoma cells impairs bone metastasis. Cancer Res. 67, 2317–24 (2007).

55. Lee, J. H. et al. Transcriptional downregulation of MHC class I and melanoma de-differentiation in resistance to PD-1 inhibition. Nat. Commun. (2020). doi:10.1038/s41467-020-15726-7

56. Zhang, Y., Feng, X. H. & Derynck, R. Smad3 and Smad4 cooperate with c-Jun/c-Fos to mediate TGF-beta-induced transcription. Nature 394, 909–13 (1998).

57. Fujii, M. et al. TGF-β synergizes with defects in the Hippo pathway to stimulate human malignant mesothelioma growth. J. Exp. Med. 209, 479–94 (2012).

58. Piersma, B., Bank, R. A. & Boersema, M. Signaling in Fibrosis: TGF-β, WNT, and YAP/TAZ Converge. Front. Med. 2, 59 (2015).

59. Ramsdale, R. et al. The transcription cofactor c-JUN mediates phenotype switching and BRAF inhibitor resistance in melanoma. Sci. Signal. 8, ra82–ra82 (2015).

60. Nallet-Staub, F. et al. Pro-Invasive Activity of the Hippo Pathway Effectors YAP and TAZ in Cutaneous Melanoma. J. Invest. Dermatol. 134, 123–132 (2014).

61. Viswanathan, V. S. et al. Dependency of a therapy-resistant state of cancer cells on a lipid peroxidase pathway. Nature 547, 453–457 (2017).

62. Antony, J., Thiery, J. P. & Huang, R. Y.-J. Epithelial-to-mesenchymal transition: Lessons from development, insights into cancer and the potential of EMT-subtype based therapeutic intervention. Phys. Biol. (2019). doi:10.1088/1478-3975/ab157a

63. Giannelli, G., Villa, E. & Lahn, M. Transforming Growth Factor-as a Therapeutic Target in Hepatocellular Carcinoma. Cancer Res. 74, 1890–1894 (2014).

64. Rodón, J. et al. Pharmacokinetic, pharmacodynamic and biomarker evaluation of transforming growth factor-β receptor I kinase inhibitor, galunisertib, in phase 1 study in patients with advanced cancer. Invest. New Drugs 33, 357–70 (2015).

65. Chakravarthy, A., Khan, L., Bensler, N. P., Bose, P. & De Carvalho, D. D. TGF-β-associated extracellular matrix genes link cancer-associated fibroblasts to immune evasion and immunotherapy failure. Nat. Commun. 9, 4692 (2018).

66. Rizos, H. et al. BRAF inhibitor resistance mechanisms in metastatic melanoma: spectrum and clinical impact. Clin. Cancer Res. An Off. J. Am. Assoc. Cancer Res. 20, 1965–1977 (2014).

67. Hugo, W. et al. Genomic and Transcriptomic Features of Response to Anti-PD-1 Therapy in Metastatic Melanoma. Cell 165, 35–44 (2016).

68. Akbani, R. et al. Genomic Classification of Cutaneous Melanoma. Cell 161, 1681–1696 (2015).

69. Joung, J. et al. Genome-scale CRISPR-Cas9 knockout and transcriptional activation screening. Nat. Protoc. 12, 828–863 (2017).

70. Gilot, D. et al. A non-coding function of TYRP1 mRNA promotes melanoma growth. Nat. Cell Biol. 19, (2017).

71. Li, W. et al. MAGeCK enables robust identification of essential genes from genome-scale CRISPR/Cas9 knockout screens. Genome Biol. 15, 554 (2014).

72. Gilot, D. et al. RNAi-Based Screening Identifies Kinases Interfering with Dioxin-Mediated Up-Regulation of CYP1A1 Activity. PLoS One 6, e18261 (2011).

73. Zhao, S., Guo, Y., Sheng, Q. & Shyr, Y. Advanced heat map and clustering analysis using heatmap3. Biomed Res. Int. 2014, 986048 (2014).

74. Anaya, J. OncoLnc: linking TCGA survival data to mRNAs, miRNAs, and lncRNAs. PeerJ Comput. Sci. 2, e67 (2016).

75. Merdrignac, A. et al. A novel transforming growth factor beta-induced long noncoding RNA promotes an inflammatory microenvironment in human intrahepatic cholangiocarcinoma. Hepatol. Commun. 2, 254–269 (2018).

76. van Zijl, F. et al. A human model of epithelial to mesenchymal transition to monitor drug efficacy in hepatocellular carcinoma progression. Mol. Cancer Ther. 10, 850–60 (2011).

77. Aibar, S. et al. SCENIC: single-cell regulatory network inference and clustering. Nat. Methods 14, 1083–1086 (2017).

