## Supplementary for "Gain-of-function CRISPR screens identify tumor-promoting genes conferring melanoma cell plasticity and therapy-resistance": Gautron et al Supplementary.pdf

#### SUPPLEMENTARY FIGURES

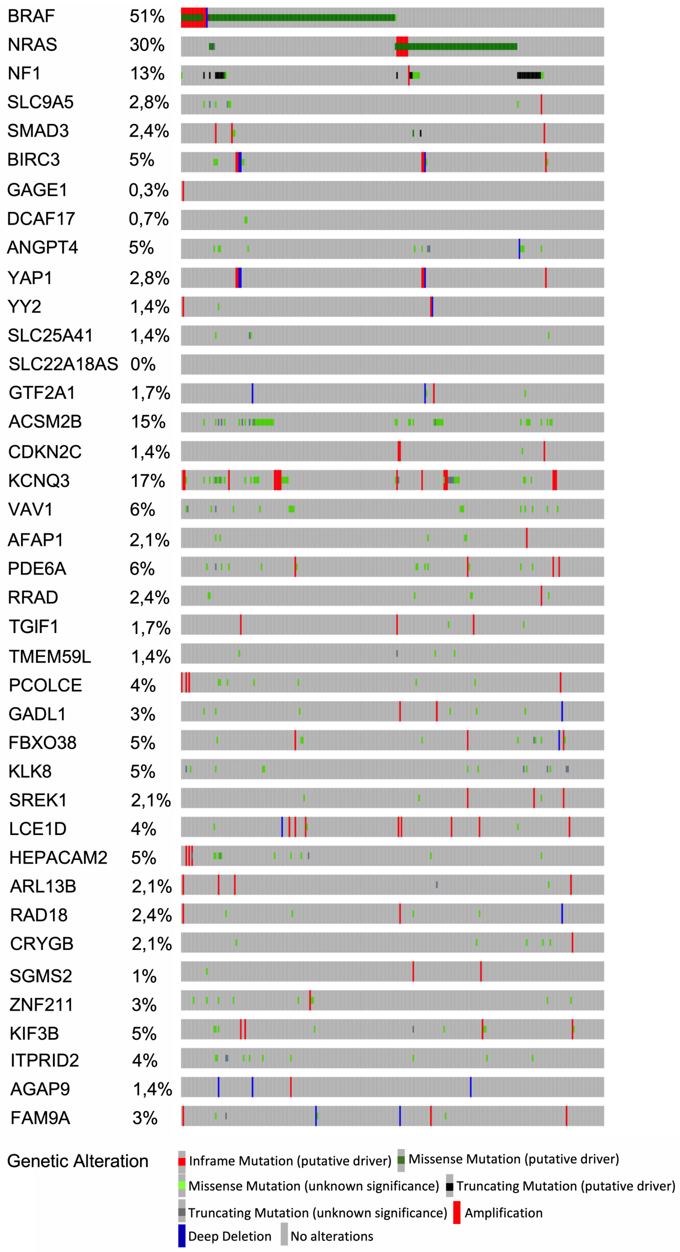

Supplementary Figure S1, related to Figure 1. Tumor-Promoting Genes are Amenable Targets for Cutaneous Melanoma.

**Supplementary Figure S1. Tumor-Promoting Genes are not frequently altered in Cutaneous Melanoma.**

Genomic alterations and tumor-promoting genes. *BRAF*, *NRAS* and *NF1* have been added to the list of tumor-promoting genes according to melanoma classification [1]. Analyses were performed using the webtool available at www.cbioportal.org.

Figure S1, related to Figure 1.

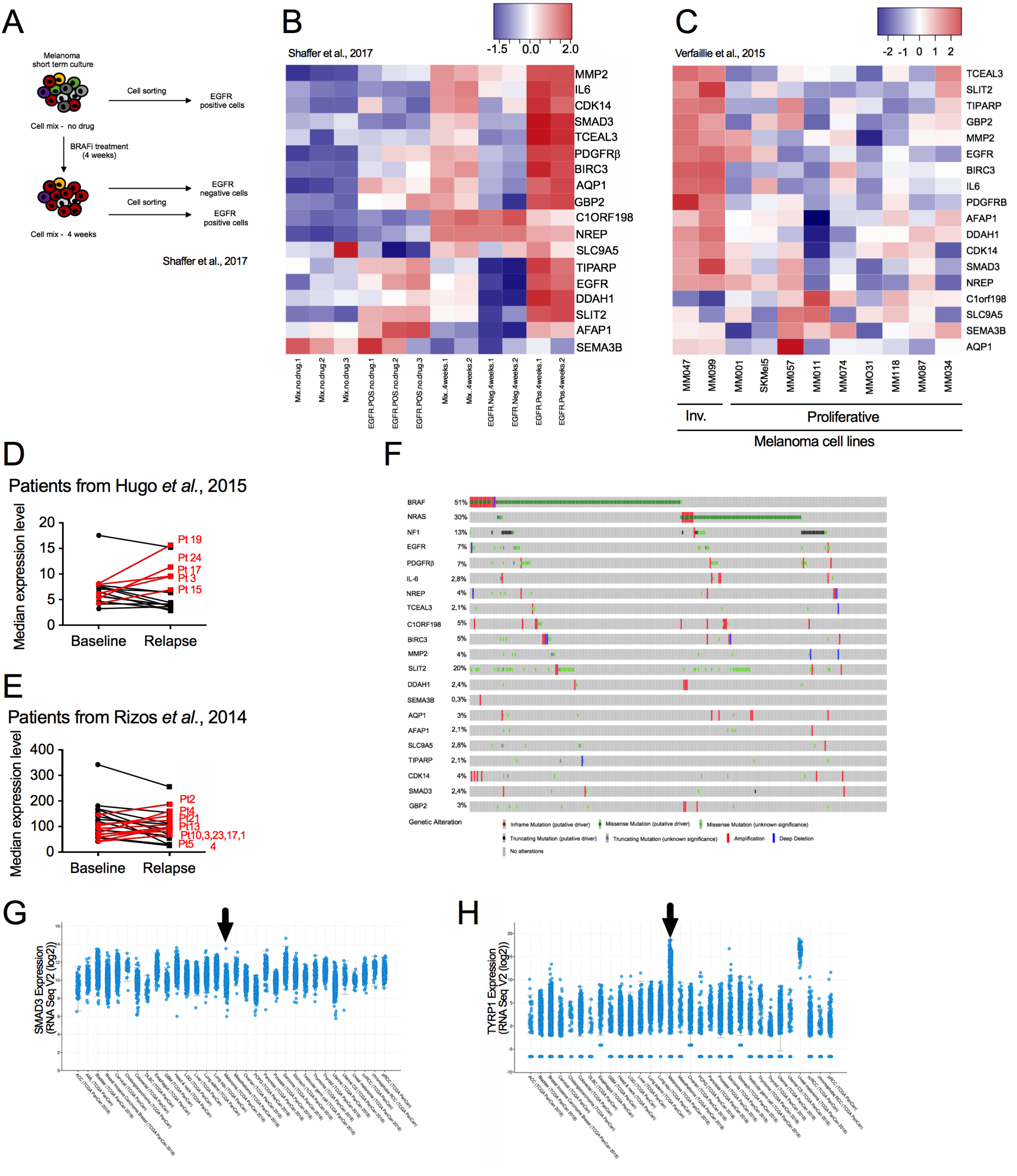

Supplementary Figure S2, related to Figure 3. BRAFi-Resistance Genes are Associated to Invasive Phenotype and Resistance to BRAFi *in vitro* & *in vivo*

**Supplementary Figure S2. BRAFi-Resistance Genes are Associated to Invasive Phenotype and Resistance to BRAFi *in vitro & in vivo.***

(A) Schematic representation of the workflow. As described in the original paper [2], fresh melanoma cells were obtained by patient tumor dissociation. Cells mix was treated with BRAFi. Cell sorting (EGFR positive and/or negative) was performed at two time points: before treatment (no drug) and after 4 weeks of treatment. Cells "mix" corresponds to the unsorted population. The EGFR-positive cells (exposed to BRAFi) are able to produce colonies.

(B) Heatmap depicting mRNA expression of BRAFi-resistance genes, using an RNA-Seq dataset obtained from BRAFi-treated melanoma cells.

(C) Heatmap illustrating the expression levels of BRAFi-resistance genes in invasive (Inv.) and proliferative melanoma cell lines (n=2 and n=9, respectively) [3].

(D) All patients from the cohort presented in Fig. 3F [4]. Expression level of our genes (median) in cutaneous melanoma biopsies before the BRAFi-treatment (baseline) and during the relapse.

(E) All patients from the cohort presented in Fig. 3G [5]. Expression level of our genes (median) in cutaneous melanoma biopsies before the BRAFi-treatment and during the relapse.

(F) Genomic alterations in BRAFi-resistance genes (from www.cbioportal.org). *BRAF*, *NRAS* and *NF1* have been added according to melanoma classification [1].

(G) *SMAD3* expression levels (RNA Seq V2 (log2) in the TCGA data set. Picture was downloaded from www.cbioportal.org. Arrows indicate cutaneous melanoma.

(H) *TYRP1* expression levels (RNA Seq V2 (log2) in the TCGA data set. Pictures was downloaded from www.cbioportal.org. Arrows indicate cutaneous melanoma. *TYRP1* is highly expressed in drug naive tumors; cutaneous and uveal melanoma in contrast to *SMAD3*.

Figure S2, related to Figure 3.

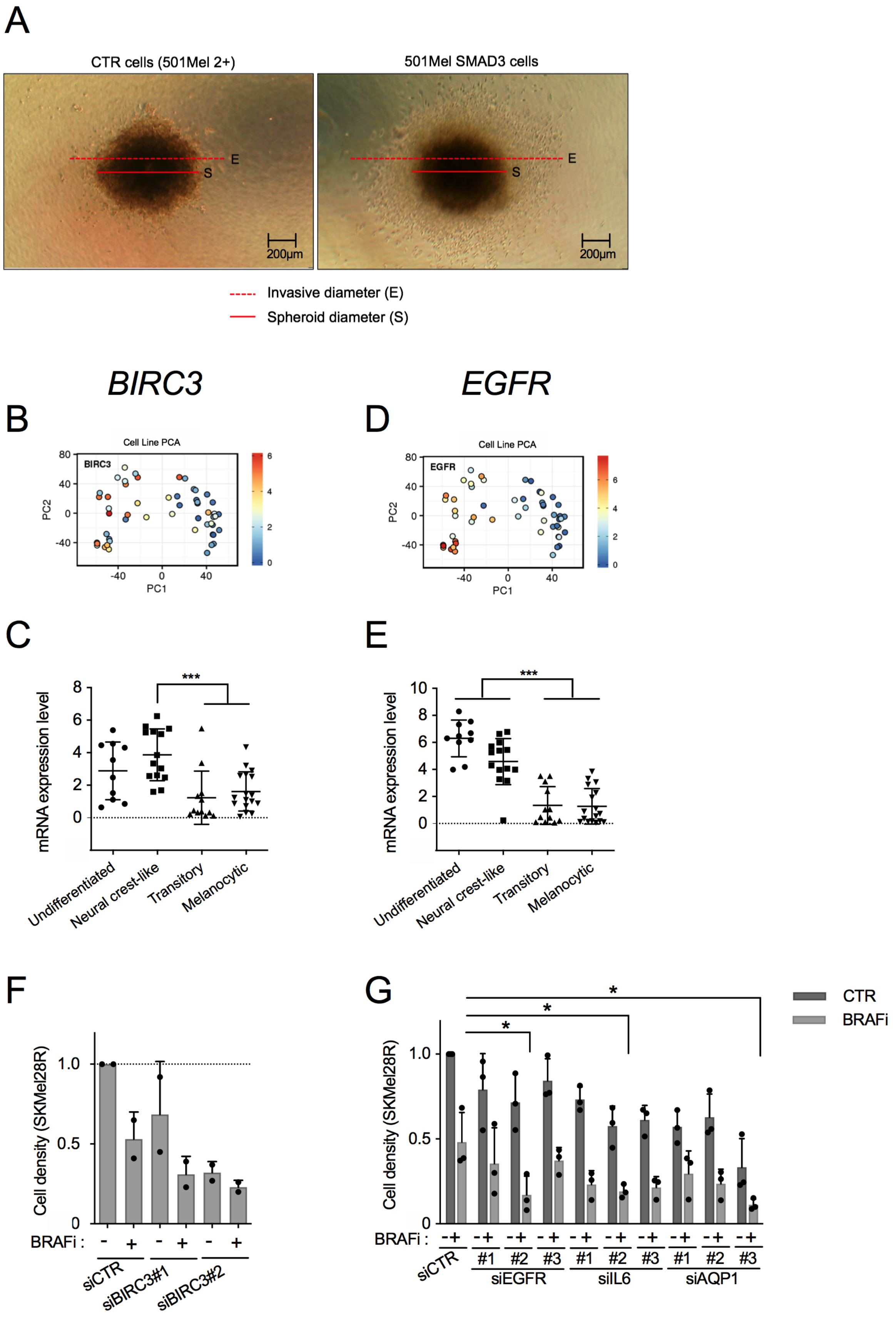

Supplementary Figure S3, related to Figure 5.
*BIRC3* and *EGFR* are Potent BRAFi-resistance Genes

**Supplementary Figure S3. *BIRC3* and *EGFR* are Potent BRAFi-resistance Genes.**

(A) Pictures illustrating the ratio (invasive diameter/spheroid diameter; E/S) used in Fig. 5E (Invasion assay, melanoma spheroids). Two cell lines are illustrated: CTR cells correspond to 501Mel cells expressing dCas9 and HSF1-p65-MS2 (named here 501Mel 2+) and the SMAD3 cells (overexpressing SMAD3).

(B) PCA analysis of *BIRC3* expression in melanoma cell lines in function of their dedifferentiation states (generated by the webtool http://systems.crump.ucla.edu/dediff/index.php). Scale: red color corresponds to a high BIRC3 expression level.

(C) *BIRC3* expression in these four subtypes of melanoma cells (undifferentiated=10; neural crest-like=14; transitory=12; melanocytic=17). Whiskers reflect mean of expression with range. Bilateral Student test (with non-equivalent variances), ***: $p<0.001$.

(D) PCA analysis of *EGFR* expression in melanoma cell lines in function of their dedifferentiation states.

(E) *EGFR* expression in these four subtypes of melanoma cells (undifferentiated=10; neural crest-like=14; transitory=12; melanocytic=17). Whiskers reflect mean of expression with range. Bilateral Student test (with non-equivalent variances), ***: $p<0.001$.

(F) *BIRC3* depletion (siRNA#1 & #2) reduced cell density and restored BRAFi (Vemurafenib) effect on BRAFi-resistant cells (SKMel28R). CTR for non-targeting siRNA. Vem for the BRAFi Vemurafenib and DMSO for dimethylsulfoxide (solvent of Vem). n=2 biologically independent experiments. Each histogram represents the mean $\pm$ s.d..

(G) Knock-down of *EGFR, IL-6 or AQP1* (siRNA#1, #2 & #3) restored BRAFi (Vemurafenib) effect on BRAFi-resistant cells (SKMel28R). n=3 biologically independent experiments are presented. Each histogram represents the mean $\pm$ s.d.. Bilateral Student test (with non-equivalent variances) *: $p<0.05$.

Figure S3, related to Figure 5.

A

Huh28 +/- TGFβ

NES=1.60
P<0.01

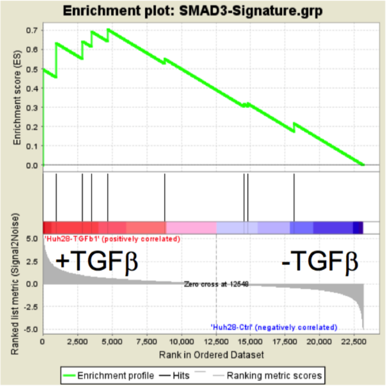

B

3sp (mesenchymal) vs 3p (epithelial)

NES=1.53
P=0.03

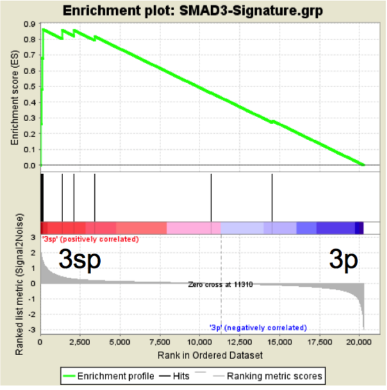

Supplementary Figure S4, related to Figure 7. SMAD3-Signature is Associated to Mesenchymal Phenotype in Liver Cancer Cell Lines

**Supplementary Figure S4. SMAD3-Signature is Associated to Mesenchymal Phenotype in Liver Cancer Cell Lines.**

(A) Gene Set Enrichment Analysis (GSEA) of the SMAD3 signature in TGF-β treated *versus* non-treated Huh-28 cholangiocarcinoma cells [6].

(B) Gene Set Enrichment Analysis of the SMAD3 signature in hepatoma cell lines : mesenchymal (3sp cells) *versus* epithelial (3p cells) [7].

Figure S4, related to Figure 7.

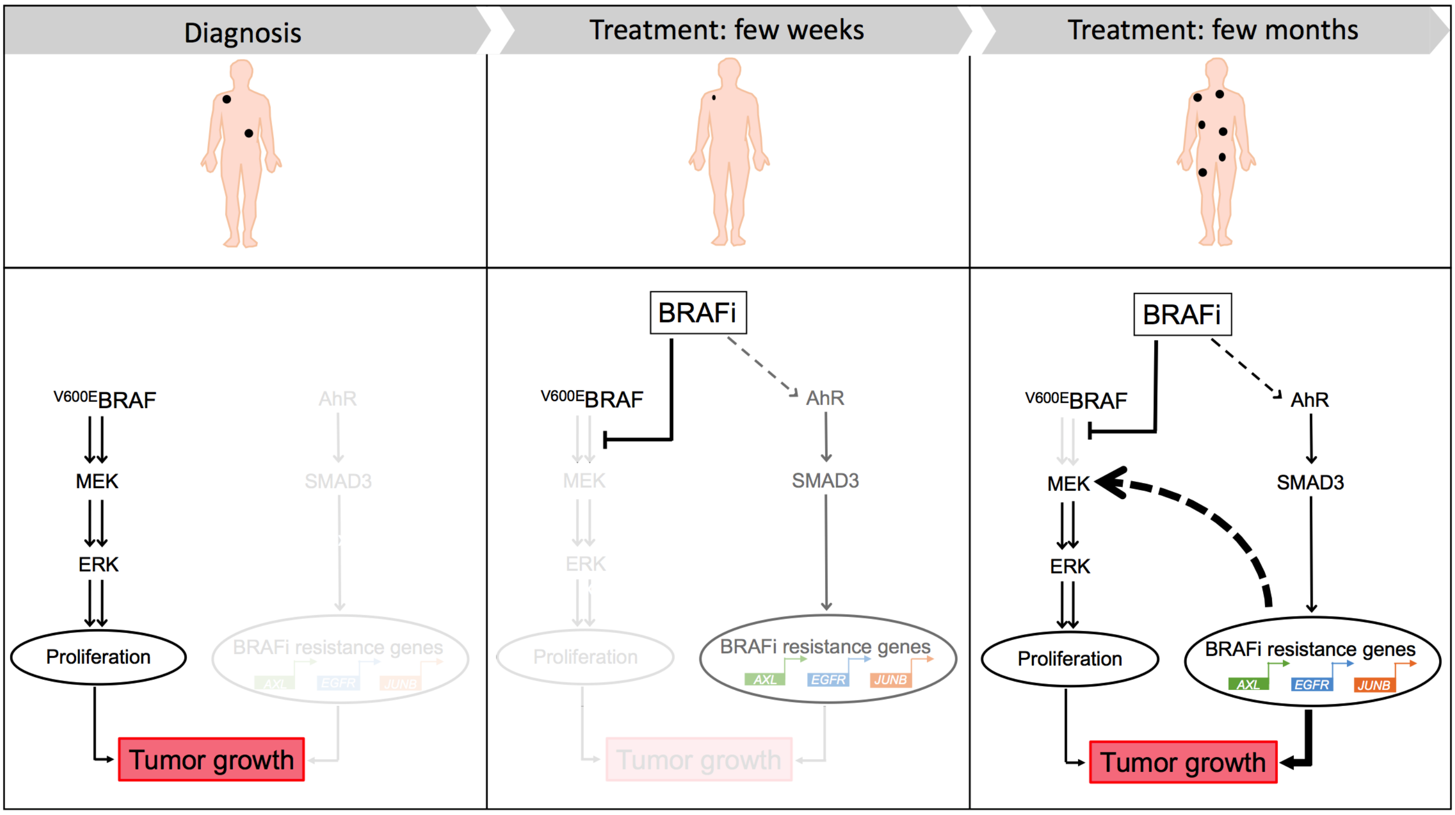

Supplementary Figure S5 An AhR-SMAD3-regulated Gene Program Promotes Therapy-resistance in Cutaneous Melanoma.

**Supplementary Figure S5. An AhR-SMAD3-regulated Gene Program Promotes Therapy-resistance in Cutaneous Melanoma.**

A scheme recapitulating the role of AhR-SMAD3 axis in BRAFi-resistance and mesenchymal transition of melanoma cells under BRAFi. V600E/K mutated BRAF constitutively activates the MEK-ERK pathway, promoting an anarchic cell proliferation and tumor growth. BRAF inhibitors (BRAFi) abrogates this deleterious activation of MAPK pathway, which is usually associated to tumor regression (few weeks). However, persister cells (BRAFi-resistant) fuel the relapse a few months later. This phenomenon is generally accompanied by a reactivation of survival pathways such as MAPK or PI3K/AKT pathways [8]. Here, we demonstrated that BRAFi resistance is associated to the AhR-SMAD3 axis. In response to BRAFi, AhR drives SMAD3 expression, which in turn transactivates potent BRAFi-resistance genes such as *AXL*, *EGFR* and *JUNB*. This reprogramming promotes BRAFi resistance and probably the reactivation of MAPK pathway. Chemical or genetic inhibition of SMAD3 decreases the expression of these BRAFi-resistance genes and restores sensitivity to BRAFi. Our work expands our understanding of the biology underlying resistance to BRAFi and highlight novel drug vulnerabilities that can be exploited to develop long-lasting antimelanoma therapies.

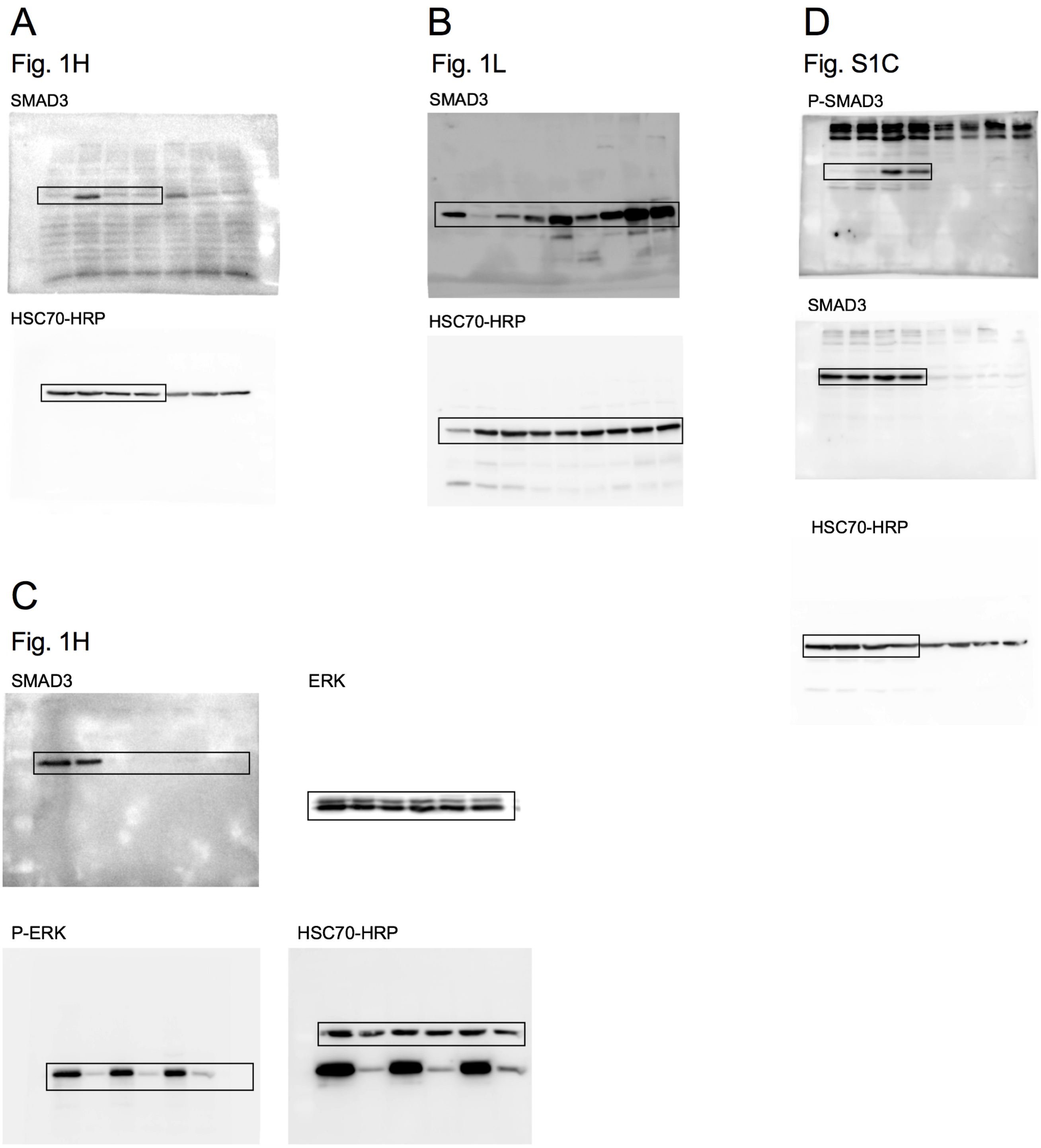

Supplementary Figure S6: Western-blot

**Supplementary Figure S6. Uncropped Western-blots**

### SUPPLEMENTARY TABLES

**Supplementary Table S1: Tumor-promoting genes**

sgRNAs counts in 501Mel cell library (DMSO 0) and in 7 tumors rising from CRISPR-SAM engineered cells (DMSO 1-7).

**Supplementary Table S2: Tumor growth curves from Fig. 1D, 1H & 4B. Raw data.**

**Supplementary Table S3: sgRNAs Occurrence in 7 tumors emerging from CRISPRa-engineered 501Mel cell library.**

Genes with sgRNA in top 100 most abundant sgRNAs among the 7 tumors developed from 501Mel cell library xenografts.

**Supplementary Table S4: Tumor growth genes and pathways annotation.**

Tumor Growth genes and Enrichr analysis (https://amp.pharm.mssm.edu/Enrichr). Rank in function of combined score. Top ten.

**Supplementary Table S5: CRISPR-SAM screen data - Normalized sgRNA counts.** Median-normalized sgRNAs counts to adjust for the effect of library sizes and read count distributions of the CRISPRa screens were assessed in 501Mel cell library (DMSO), BRAFi-resistant cells (Vem for Vemurafenib) and PLX8394 (PB for Paradox Breaker). In each condition, sgRNA were sequenced from two independent experiments and reads were pooled before counting and normalization. Raw sequencing data of the CRISPRa screen (fastq files) are publicly available in ArrayExpress under the accession E-MTAB-8595.

**Supplementary Table S6: Number of sgRNAs per gene enriched in BRAFi-resistant cells.** To test whether sgRNA abundance differs significantly between BRAFi and DMSO arms, sgRNAs median-normalized counts (Table S5) were compared using a negative binomial model implemented in the MAGeCK algorithm. sgRNAs with a false discovery rate <= 0.05 were conserved and the number of significant sgRNAs/gene are reported for Vem (Vemurafenib), PLX8394 (PB, Paradox Breaker) and the combination of the two BRAFi screens (PB+Vem; PBV).

**Supplementary Table S7: sgRNAs Occurrence in 7 tumors emerging from BRAFi-persister cells (CRISPRa-engineered 501Mel cell library exposed to BRAFi).**

**Supplementary Table S8: SMAD3-Signature**

List of genes creating the SMAD3-signature (established from Figure 7A).

**Supplementary Table S9: References for antibodies, siRNA and sequences of primers and sgRNAs.**

**Supplementary Table S10: Statistics data source**

### Supplementary Methods

#### CRISPR-SAM Screen

**Protocol adapted from Joung J.** et al., Nat Protoc. 2017 (doi:10.1038).

**ACKNOWLEDGEMENTS**
The authors are grateful to Feng Zhang for providing the Human CRISPR 3-plasmid activation pooled library (SAM) (Addgene). Human CRISPR Activation Library (Pooled library) – Addgene #1000000057
This library consists of three components which are all provided:

1. A nucleolytically inactive Cas9-VP64 fusion (Addgene plasmid # 61425). Blasticidin resistance.
2. A gRNA incorporating two MS2 RNA aptamers at the tetraloop and stem-loop 2 (present in the libraries)
3. The MS2-P65-HSF1 plasmid which expresses the activation helper protein (Addgene plasmid #89308). Hygromycin resistance.

#### Generation of 501Mel 2+ cells (dCas9 and MS2-P65-HSF1)

501Mel cells (200,000 cells per well (MW6)) have been co-infected overnight with viral supernatants (multiplicity of infection, MOI~0.2) produced as detailed below:

**D0:** plating of **two T25** of HEK293T (DMEM + 10% FCS + 1% PS) at about 40% confluence.

**D+1:** at about 50-60% confluence, perform the transfection according to the following conditions:

For one T25: 3.2 µg lentiviral plasmid + 1.1µg pVSV-G + 2.1µg psPAX2 into 180µL of OptiMem (tube A) and 16 µL Lipofectamine 2000 into 180µL (tube B).

pool tube A into tube B: mix with your finger (10x) and incubate 15 min (RT).
Distribute on cells (containing 3 mL DMEM + 10% FCS + 1% PS).

**5h after transfection**, change the cell medium. Add the minimum volume (3 ml/flask T25).
**D+3:** collect the infectious medium and centrifuge it to discard cells and cell debris (1300rpm, 3min, RT). Carefully collect the supernatant and filter it on PVDF filter 0.45µm with a low pressure.

Cells have been exposed to infectious media in presence of polybrene (8µg/mL) overnight. The MOI is ~0.2. Conventionally, the viral titer obtained in our conditions is ~1 x$10^6$ TU/mL (TU for transduction unit).

After 5 days of antibiotic selection (Blasticidin (2 µg/mL) and Hygromycin B (200 µg/mL)), expression levels of dCas9 and MS2-p65-HSF1 were evaluated by RT-qPCR. Primer sequences are available in Gautron et al., Nat. Comms 2020). These cells correspond to the "501Mel 2+ cells".

### Amplification of the plasmid library lentiSAMv2 (cat. no. 1000000057)

#### I - Preparation of LB medium

##### LB Lennox to prepare plate:

References:
- Petri (Ø90mm h14,2mm) – VWR #391-0455.
- Squared Petri : NUNC for culture (245 x 245 x 25 mm; 500cm$^2$) – Dutscher #055064

| Powder | gram | Ref. | Batch |
|---|---|---|---|
| Tryptone | 10g | BD #211705 | 5027930 |
| Yeast Extract | 5g | BD #212750 | 3326451 |
| NaCl | 5g | Sigma #S7653 | SZBE2100V |
| Agar | 15g | BD #214010 | 5232955 |

Fill to 1 liter with $H_2O$ milliQ. **Adjust pH to 7.0**.
Add a bar magnet.
Autoclave the medium: short cycle.
Allow the medium to return to a temperature of about 55 ° C in a water bath to avoid caking.
Add the antibiotic (Ampicillin 100µg/ml), homogenize on a magnetic plate (not by turning over to avoid bubbles).
Pour the petris. Wait for the setting in mass (several hours). Store at + 4 ° C. Protect plates with plastic food film and plastic bag. Check them one week later for sterility.

##### LB Miller to collect the colonies on the boxes:

| Powder | gram | Ref. | Batch |
|---|---|---|---|
| Tryptone | 10g | BD #211705 | 5027930 |
| Yeast Extract | 5g | BD #212750 | 3326451 |
| NaCl | 10g | Sigma #S7653 | SZBE2100V |

Fill to 1 liter with $H_2O$ milliQ. **Adjust pH to 7.0.**
Autoclave the medium: short cycle.
Allow the medium to cool, and store at + 4 ° C.

#### II –Bank Amplification

Amplification of the SAM library is performed by electroporation of electrocompetent bacteria.

References:

- Electroporator : Bio-Rad Micro Pulser (Cat. #165-2100)
- Electroporation vats : Bio-Rad Gene pulser cuvette 0.1cm 50 pK (Cat. #1652086)
- Bacteria : Lucigen E cloni. 10G Elite (Cat. #60051-2) ; batch n°11514
- Lucigen pUC19 DNA -10pg/µL (Pt#F92078-1) ; batch n°11158
- Lucigen Recovery medium (Cat. #80026-1) ; batch n°11390
- Human CRISPR Activation Library (Pooled library) – Addgene #1000000057
- Maxiprep : NucleoBond Xtra Maxi (Macherey-Nagel ; ref : 740414.10)
- Conical centrifugation pots 250mL (Corning ; ref : 430776)

**NOTE:** Before carrying out the amplification of the library, the transformation efficiency must be tested using the equipment listed above.

- **Electroporation trial:**

25 µL of electrocompetent bacteria were transformed with 10 pg of pUC19 (available with bacteria).
Electroporation was performed with the "bacteria" ("standard") mode of the electroporator.
After the electric shock, the bacteria were taken a few minutes later (the time to go into bacteriology room) by 2mL of "recovery medium".
Isolation was performed on a Ø90mm petri plate (LB Lennox) from 10µL of the diluted bacterial solution above.

After an overnight incubation at 37 ° C (box turned), 100 colonies were counted on the plate.
100 colonies for 10µl from 2mL (recovery medium). So, for 10 pg used, 20,000 colonies were obtained. Thus, for 1µg, we would obtained 2 x$10^9$cfu / µg. Efficiency = 2x$10^9$ cfu / µg.
Minimum recommended efficiency for the amplification of the library = 1 x$10^9$cfu / µg.

**Note:** Size of pUC19(2,7kb) is different to lenti sgRNA MS2 (9.9kb). So, the efficiency will be less efficient with the lentiviral plasmid.

- **Plasmid library Amplification:**

To amplify the plasmid library, it is necessary to have poured (LB Lennox) at least 8 squared boxes NUNC (245 * 245 * 25 mm) and 1 petri dish Ø90mm.
**Note**: It is advisable to prepare much more to overcome various problems (agar damaged, etc.). It is therefore advisable to prepare at least 12 squared boxes and 3 boxes of petri dishes. Before beginning the transformation, incubate these dishes, the lid on top, in the incubator at 37 ° C for about 2 hours. Keep them at 37°C during the transformation.

The following steps are performed around flame (bunsen burner) on a bench near the electroporator.

1µL of the bank (eq 50ng) is added on 25µL of bacteria. The mix is gently homogenized by 2 go/return movements into the tip of the pipette 200 before transfer into the electroporation cell.
**Note:** Be careful, do not make a bubble when you put the mix in the tank! Electroporation was performed with the "bacteria" ("standard") mode of the electroporator. Then, as soon as possible, 975µL of "sterile recovery medium" is gently added to the tank, **without going back and forth** but trying to create a slight current in the bottom of the tank during the addition. Then transfer everything into a sterile 2mL Eppendorf that is placed on ice.

**This step is carried out 8 times (1 µl of plasmid library on 8 different tubes of bacteria).** To facilitate the handling, it is easier to work with somebody: one who deals with the "mix" then transfer into the tank and at the end of the addition of the medium, and another one who performs the electroporation. This allows to be faster.

Once the 8 repetitions are done, add 1mL of "recovery medium" in all the tubes.
**Note**: Also prepare a 2mL Eppendorf tube with 1mL of "recovery medium" for calculating the transformation efficiency.

Climb bacteriology room. Incubate the 8 tubes at 37°C for **1 hour.**

Under laminar flow: Pool the 8 tubes (16mL in total) in a sterile 50mL Falcon.

**Note**: Take 20µL that is added to the tube containing 1mL of "recovery medium" prepared previously. Spread 100µL of this solution on the Ø90mm petri dish and incubate bottom lid at 37°C overnight. It will be used to calculate the transformation efficiency of the current experiment.

Always Under laminar flow, **spread 2 mL of the bacterial pool** (Falcon 50mL) **on each square box** using a "blue rake". Let "drink" these boxes at 37 ° C, cover up, for ~1h before turning and incubate on the night.

Before recovering the bacteria from the squared boxes, it is necessary to validate the efficiency of transformation thanks to the petri dish prepared previously.

On this box, we got ~300 colonies on 1/8th of the box. That is about 2400 colonies in total. So, 2400 colonies for 100µL. These 100µL correspond to 1/10th of the 2mL solution containing 20µL of bacteria from the 16mL spread in total. So total colony number = 2400 * 10 * 8000 = 1.92 x$10^8$colonies.
It is recommended a minimum of 7 x$10^7$colonies (eq 100X colonies per guide).

**NB: think about weighing the conical pots that will be used to pack the bacteria.**

After removing the plates from the incubator, they are scraped 2 by 2. Indeed, to recover the bacteria, 10mL of LB Miller medium are added to each box, which are scraped with a small cell scraper. The detached bacteria are then transferred to a conical pot for centrifugation. A second phase of scraping is carried out on each box with 10ml of medium, and the bacteria recovered in this same plot. Two pots are used: 4 plates / pot.

Once the assembly is complete, the bacteria are centrifuged for 30 min - 3000 rpm - 4 ° C. The supernatant is carefully removed by inversion and the pots and the pellets are weighed. **Otherwise the LB distorts the weighing.** Pot n ° 1: 33,87g - Pot n ° 2: 33,71g. Therefore, weight of the pellets: No. 1 pellet = 4.22 g - No. 2 pellet = 3.86 g**.**

The plasmids of the library are purified using a kit of Maxiprep, at the rate of **a maxiprep for about 0.75g of bacteria. Here we used 11 maxipreps**. Plasmids were checked on BET gel and nanodrop.
Aliquots were stored at -80°C.

### From Plasmids to Lentivirus

#### I –Lentivirus Production

**D0:** prepare **twelve T225** of HEK293T cells (DMEM + 10% FCS + 1% PS) at about 40% confluence. (6 full flasks T225 full to generate 12 **flasks T225 containing** $30x10^6$ HEK293T cells).

**D+1:** at about 50-60% confluence, perform the transfection according to the following conditions:

For one T225: 10µg sgRNA library + 10µg pVSV-G + 15µg psPAX2 and 90µL Lipofectamine 2000. So, for each virus production: 120µg VSV-G and 180µg PAX2 and 120µg library)

For **12**flasks T225:

| | | | |
|---|---|---|---|
| Tube A: pVSVG | 1172 ng/µL | 120 µg | 102 µL |
| Tube A: psPAX2 | 934 ng/µL | 180 µg | 193 µL |
| Tube A: SAM bank | 2478 ng/µL | 120 µg | 48µL |
| Tube A: OPTI-MEM | 16,2 mL | | |
| Tube B : Lipo2000 | 1080 µL | | |
| Tube B: OPTI-MEM | 16,2 mL | | |

Mix tube A into tube B: mix with your fingers (10x) and incubate 15 min (RT).
Distribute 2.8 mL per T225 (containing 25mL DMEM + 10% FCS + 1% PS).

**5h after transfection**, change the cell medium. Add the minimum volume (25 mL/flask).
**D+3**: Collect the infectious media and centrifuge them (1300rpm, 3min, RT) in falcon 50mL.
Collect the supernatants (25mL) and filter them **with a low pressure** (50mL syringe) on a **0.45µm PVDF** filter (**NOT cellulose filter**). **ONE filter per T225** (25mL).
Pool supernatants and homogenize gently. **To use immediately (about 300mL). If you store this viral supernatants, ~50% of virus is destroyed. Keep in mind for MOI calculation.**

**NOTE : D+2:** 4 x T225 containing $10x10^6$ 501Mel2+ were prepared cells (RPMI + 10% FCS + 1% PS). Six petris must also be prepared (Ø100mm) with $1x10^6$ of 501Mel2+ cells/petri for the MOI calculation.

• **<u>The Multiplicity of infection sought is 0.2</u>**. From the observations of the first experiment, we used a volume of 40mL of virus (0.22x$10^6$ TU/mL) for 40M of cells. (40mL x 0.22x$10^6$ TU/mL = 8.8 x$10^6$ TU for 40x$10^6$ cells. MOI=0.22. **It is important to keep in mind that at an MOI 0.3 or less, greater than 95% of infected cells are predicted to have a single integration and is therefore recommended for pooled screening.**
8.8 x$10^6$ TU and the library contains 70290 sgRNA (guides). In this case we will obtain ~125 infected cells expressing the same guide). So, 10mL of viral supernatant have been added per flask with polybrene (4µg/µL final). Infection overnight.

#### II –MOI Calculation (post-infection)

The infection should be made at a MOI of 0.2. To test the MOI, carry out an infection of 501Mel2+ in P100 (1M of 25% confluent cells) with 6 different dilutions of lentivirus :100µL/petri (100th, 200th, 500th, 1000th, 2000th and without virus).
First prepare 500µL of viral supernatant diluted 1: 100 (5 µL + 495 µL medium)

| Dilution | **Infectious medium** | Volume medium |
|---|---|---|
| 1/100 | 100 µL | / |
| 1/200 | 50 µL | 50 µL |
| 1/500 | 20 µL | 80 µL |
| 1/1000 | 10 µL | 90 µL |
| 1/2000 | 5 µL | 95 µL |

If you have time, it is interesting to test additional dilution 1/50, 1/20, etc.

After 3 to 6 days of antibiotic selection (zeocin at 600 µg/mL). Count surviving colonies after staining with methylene blue (Gilot D. et al., Nat Cell Biol 2017). **NO colony in petri without virus (negative control) at the end of antibiotic selection.**
For example, 110 colonies were counted in the 1/200 dilution, which indicates 22,000 TU/100µL of this dilution (1/200). Thus, our viral supernatant contained 0.22x$10^6$ TU/mL. Similar results were obtained with the other dilutions (41 for 1/500 et 20 for 1/1000). We estimated the titer ~0.2x$10^6$ TU/mL.

#### <u>III–Cell Selection</u>

**D+1** (24 h post infection): count the cells and split them in several T225 at the rate of 7.5 x$10^6$ cells per T225 (low density seeding) to obtain a better and faster antibiotic selection. Knowing that we seeded 10M by T225 two days ago, we consider that we will obtain ~25M cell by T225, so 100 x$10^6$ cells per condition. **Provide 14 T225/condition.**

**3h after the plating** (cells are adherent), **apply the antibiotic selection** (here : zeocin (300 µg/mL then 600 µg/mL).

| Day | Amount | Volume (medium) | Volume Zeocin |
|---|---|---|---|
| D+1 | 300µg/mL | 980mL | 2940 µL |

| | | | | |
|---|---|---|---|---|
| D+3 | 600µg/mL | 980mL | 5880 µL | |
| D+5 =end of selection | 0µg/mL | 980mL | 0 | If selection is not complete, expose cells to 600µg/mL |
| Total | | 2940 mL | **8820** µL | Or **14700**µL |

**D+7:** Split the cells in T225 at ~6-7 M cell/T225.
**As soon as possible, make a pellet of at least 36 x10$^6$ cells and make freezing tubes with the same amount.**
For the screen with this library, it is necessary to work with at least 500 cells per sgRNA (500x70,290 = 35,145,000 cells). It is therefore necessary to keep at least 36x10$^6$ cells for the experiments. We used 40x10$^6$ cells for each experimental condition.

**Note:** In our manuscript (Fig.1, Gautron A. et al. 2020), we estimated the coverage (>95%), the sgRNA distribution (mean 550) and the correlation between replicates (Control (DMSO): r=0.86 and for BRAFi: r=0.95).

#### From Cell Bank to Resistant Cells

**Note:** If you start from already infected and frozen cells, **you have to do a centrifugation to eliminate the medium + DMSO, otherwise the cells do not adhere well**.

**D+7:** Resuspend the cells and separate them in different conditions according to the treatments. Here DMSO and BRAFi. Therefore, 40x10$^6$ cells * 3 = 120 x10$^6$ cells. Cells were split to control (DMSO= solvent of BRAFi) and drug treatment arms (BRAFi; PLX4032 (2µM, Selleckchem) or Paradox Breaker (PLX8394, 2µM, MedChemExpress)). **During the treatment (14 days), media were renewed every two days** in order to eliminate dead cells and to ensure a potent BRAFi pressure.

**D+21:** After 14 days of treatment, make a pellet of each condition. Cells were pelleted by centrifugation, resuspended in PBS, and frozen promptly for genomic DNA isolation to identify sgRNA sequences. Pellets could be stored (-80°C).

#### From Cell Pellets to Guide Identification

For this part, we used the protocol published by Julia Joung (Nature Protoc, 2017).
Genomic DNAs from cell pellets and tumors (>400mg) were extracted using the Zymo Research Quick-gDNA MidiPrep according to the manufacturer's protocol. PCR amplifications and quality controls have been done as described by Zhang lab (Joung et al., 2017).

**<u>I –Library</u>**

- Genomic DNA was extracted using the Zymo Quick-gDNA midi kit (Zymo Research). One pellet split on 4 columns (max 10x10$^6$ cells/column). We obtained at least 250µg/pellet.

- DNA Quantification has been made with Nanodrop and BioAnalyzer.
- PCR have been performed using the NEBNextHigh Fidelity 2X Master Mix. 32 cycles/single-step reaction.

For one sample, library is made using 10 forward primers (diversity) for one Reverse primer (Reverse primer used for multiplexing). Eight PCR per primer couple. So, we generate 80 wells per sample.

2.25µg/well (PCR).

For another sample, use the same 10 forward primers but change the Reverse primer (cf supplementary Table Konermann et al., Nature 2015).

Purify pooled PCR products using Zymo spin V with reservoir

PCR products have been separated on agarose Gel (2%) to eliminate nucleotides and non-desired amplicons. **Note: check the bubble product (generated by PCR) with the BioAnalyser and on agarose gel. Only one PCR product should be detected and purified.**

**Illustration: BioAnalyser profile of PCR-products before gel migration. Expected size (bank)~280bp. Bubble product >400bp.**

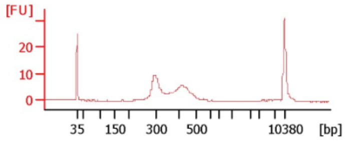

PCR products have been Gel extracted using Zymoclean Gel DNA Recovery Kit - Uncapped columns

The library is sequenced on HiSeq (35 million reads passing filter per library).

#### II –Sequencing

Sequencing was performed by the Human & Environmental Genomics core facility of Rennes on a HiSeq 1500 (Rapid SBS kit v2 1x100 cycles, Illumina). Base Calling was performed with Illumina's CASAVA pipeline (Version 1.8).

#### III –SgRNA Enrichment Analysis

Data processing was conducted using the MAGeCK v0.5.6 software. Briefly, read counts from different samples are first median-normalized to adjust for the effect of library sizes and read count distributions (mageck count with option: --norm-method median). Then, in an approach similar to those used for differential RNA-Seq analysis, the variance of read counts is estimated by sharing information across features and a negative binomial model is used to test whether sgRNA abundance differs significantly between the treatment conditions and the DMSO control. Positively or negatively selected sgRNA are ranked according to adjusted P-values (false discovery rate) and gene log fold changes computed with the modified robust ranking aggregation algorithm implemented in MAGeCK (mageck test with options: --norm-method median, --gene-lfc-method alphamedian, --adjus

### SUPPLEMENTARY REFERENCES

1. Akbani, R. *et al.* Genomic Classification of Cutaneous Melanoma. *Cell* **161**, 1681–1696 (2015).
2. Shaffer, S. M. *et al.* Rare cell variability and drug-induced reprogramming as a mode of cancer drug resistance. *Nature* **546**, 431–435 (2017).
3. Verfaillie, A. *et al.* Decoding the regulatory landscape of melanoma reveals TEADS as regulators of the invasive cell state. *Nat. Commun.* **6**, 6683 (2015).
4. Hugo, W. *et al.* Non-genomic and Immune Evolution of Melanoma Acquiring MAPKi Resistance. *Cell* **162**, 1271–1285 (2015).
5. Rizos, H. *et al.* BRAF inhibitor resistance mechanisms in metastatic melanoma: spectrum and clinical impact. *Clin. Cancer Res. An Off. J. Am. Assoc. Cancer Res.* **20**, 1965–1977 (2014).
6. Merdrignac, A. *et al.* A novel transforming growth factor beta-induced long noncoding RNA promotes an inflammatory microenvironment in human intrahepatic cholangiocarcinoma. *Hepatol. Commun.* **2**, 254–269 (2018).
7. van Zijl, F. *et al.* A human model of epithelial to mesenchymal transition to monitor drug efficacy in hepatocellular carcinoma progression. *Mol. Cancer Ther.* **10**, 850–60 (2011).
8. Welsh, S. J., Rizos, H., Scolyer, R. A. & Long, G. V. Resistance to combination BRAF and MEK inhibition in metastatic melanoma: Where to next? *Eur. J. Cancer* **62**, 76–85 (2016).
